# Multi-compartment immune and tumor cell reprogramming by LNP-IFNα2 overcomes colon cancer immunotherapy resistance

**DOI:** 10.64898/2026.06.26.734809

**Authors:** Zainab Tiamiyu, Patrick Czabala, Jennifer Worthy, Luis Velasquez Zarate, Mei Zheng, Dakota B. Poschel, Kendra Fick, Dafeng Yang, Henry R. Monnig, Richa Rashmi, Sergei Bombin, Priscilla S. Redd, Kebin Liu

**Affiliations:** Department of Biochemistry and Molecular Biology, Medical College of Georgia, Augusta University, Augusta, GA; VA Augusta Health Care System, Augusta, GA; Department of Pathology, Medical College of Georgia, Augusta University, Augusta, GA; Georgia Cancer Center, Medical College of Georgia, Augusta University, Augusta, GA; CheMedImmune Inc., Augusta, GA. USA

**Author notes:** Correspondence: Priscilla S. Redd, or Kebin Liu. 1410 Laney Walker Blvd, Augusta, GA 30912. USA. Equal contributions (Zainab Tiamiyu, Patrick Czabala).

## Abstract

Approximately 85-90% of colorectal cancers (CRC) are microsatellite stable and resist immune checkpoint inhibitors (ICI). The recent success of intismeran, a lipid nanoparticle (LNP)-delivered mRNA encoding patient-specific tumor antigens, enhances pembrolizumab responses, showing that LNP-delivered nucleic acids can render ICI-resistant tumors responsive by supplying what the tumor microenvironment lacks. Because an IFN-responsive phenotype predicts CRC response to checkpoint blockade and tumor cell PD-L1 represses IFN-I, we asked whether delivering the missing cytokine, rather than antigen, achieves the same conversion. Here we show that PD-1 blockade failed even when initiated before tumor seeding, yet deleting tumor cell PD-L1 abolished colon cancer lung metastasis, implicating a tumor-intrinsic PD-L1 function that antibody blockade does not neutralize. An LNP-encapsulated IFNα2-encoding nanoplasmid (LNP-IFNα2) transfected tumor cells within lung metastases, converting them into a self-sustaining and tumor-restricted IFNα2 source. LNP-IFNα2 suppressed colon cancer lung metastasis in syngeneic and humanized mouse models and sensitized these ICI-resistant tumors to PD-1 blockade. Efficacy improved further with neutralization of co-induced IL6. scRNA-Seq revealed multi-compartment reprogramming: SPP1^+^ macrophages acquired apoptotic signatures as IFN-responsive monocytes replaced them, progenitor-exhausted T cells exited quiescence and expanded, and tumor cells exited a high-cycling state, gained antigen presentation, and shifted away from cuproptosis resistance. The treated microenvironment transcriptionally recapitulated pembrolizumab-responder signatures, positioning LNP-IFNα2 as an off-the-shelf nanomedicine for ICI-resistant CRC.

## INTRODUCTION

Immune checkpoint inhibitor (ICI) immunotherapy has been approved for colorectal cancer (CRC) patients harboring mismatch repair-deficient (dMMR) or microsatellite instability-high (MSI-H) tumors^1,2^. However, fewer than 50% of dMMR/MSI CRC patients respond durably^1,3^, and, critically, approximately 85-90% of CRC cases are proficient mismatch repair (pMMR) tumors that are broadly non-responsive to ICI therapy^4,5^. No effective strategy exists to convert checkpoint-resistant CRC into an immunotherapy-responsive state. The recent clinical success of intismeran, a lipid nanoparticle (LNP)-encapsulated synthetic mRNA encoding a concatemer of patient-specific tumor antigens, in enhancing responses to pembrolizumab highlights the potential of LNP-based nucleic acid therapies to overcome resistance to immune checkpoint inhibitor (ICI) immunotherapy in human cancers^6,7^.

Interferon (IFN) is an essential component of host cancer immune surveillance^8–19^. A large body of evidence implicates a defective IFN response in cancer patients as a key determinant of ICI resistance^20,21^. Loss-of-function mutations in JAK1 and JAK2, key transducers of IFN receptor signaling, confer both primary and acquired resistance to ICI immunotherapy^20–27^. Conversely, a constitutive interferon-high immunophenotype has recently been defined as a robust predictor of ICI response specifically in CRC^5^, reinforcing IFN signaling as a central axis of antitumor immune competence. However, IFN-I expression is often repressed in the tumor microenvironment by the PD-(L)1 axis^28^. Beyond the canonical engagement of PD-1 on cytotoxic T lymphocytes (CTLs) to inhibit their activation^29–31^, tumor cell PD-L1 also engages PD-1 expressed on myeloid cells, an interaction we have previously shown to repress IFN-I expression to downregulate CXCL9 expression to impair CTL recruitment to lung metastases^28^. Loss of IFN-I expression thus emerges as a critical and proximal consequence of PD-L1 immune suppression within the metastatic niche, providing a mechanistic rationale for directly restoring IFN-I rather than relying on ICI alone to relieve this suppression in a TME that has already silenced the upstream immune IFN signal^19,32^.

Restoring IFN-I in the tumor is fundamentally a delivery problem. Recombinant IFNα is an FDA-approved anticancer agent with clinical activity in melanoma and renal cell carcinoma^33–35^, but its short half-life necessitates frequent high doses, resulting in substantial toxicity^36,37^. Pegylation extends half-life and reduces dosing frequency but does not significantly reduce overall toxicity^38–40^. Engineered alternatives, including tumor protease-activated IFN prodrugs and antibody-cytokine fusions, improve tumor targeting but remain systemically administered recombinant proteins whose exposure is governed by plasma pharmacokinetics rather than by tissue localization^41,42^. A first-in-human phase 1/2a trial in glioblastoma used genetically engineered autologous hematopoietic stem cells to achieve tumor-restricted, myeloid cell-mediated IFNα2 gene therapy, demonstrating feasibility and immune microenvironment reprogramming without systemic toxicity^43^. While this establishes clinical proof-of-concept for localized IFNα2 gene therapy, it requires ex vivo lentiviral transduction and myeloablative conditioning, a bespoke cell product rather than an off-the-shelf agent. Nucleic acid delivery offers an off-the-shelf route to the same end: nanoparticle-encapsulated mRNAs encoding a multiplexed cytokine cocktail that includes IFNβ drive local cytokine expression and curative responses in autochthonous pancreatic cancer models after systemic administration, without dose-limiting toxicity^44^. That approach depends on transient mRNA expression and on combining five cytokines, as no single cytokine mRNA was sufficient, leaving open whether one durably expressed type I IFN payload can achieve the conversion. Achieving spatially confined IFNα activity therefore requires a carrier that concentrates the therapeutic payload in the metastatic niche and converts the tumor itself into the production site.

Lipid nanoparticles (LNPs) provide such a carrier. Cationic DOTAP-cholesterol LNPs exploit this principle: their net positive surface charge drives first-pass retention and preferential accumulation in tumor-bearing lung^45^, and rapidly dividing tumor cells take up and express nanoparticle-delivered plasmid DNA more efficiently than surrounding slowly dividing normal cells^46–49^. DOTAP-cholesterol-plasmid formulations have delivered tumor suppressor transgenes to lung tumors in patients with a 10-25-fold enrichment of transgene product in tumor tissue over circulating lymphocytes and without dose-limiting toxicity, and are currently in clinical testing (NCT05703971 and NCT04486833)^46^.

We developed DOTAP-cholesterol LNPs encapsulating an IFNα2-encoding nanoplasmid (LNP-IFNα2) and mRNA, and demonstrated the preferential accumulation in tumor-bearing lungs and suppression of breast and melanoma metastases^50^. Here, we advance from therapeutic efficacy to defining the underlying mechanisms and therapeutic sensitization in ICI-resistant mouse and humanized CRC lung metastasis models. We determined that tumor cells are the transfected compartment, converting lung metastases into an autonomous IFNα2 source via an early acute cytokine phase and a durable transgene phase, This sensitizes checkpoint-refractory colon cancer lung metastases to PD-1 blockade through IFNAR1-dependent TME reprogramming: SPP1^+^ macrophages acquired apoptotic signatures as IFN-responsive monocytes replaced them, progenitor-exhausted T cells exited quiescence and expanded, and tumor cells lost their high-cycling state, gained antigen presentation, and shifted away from a cuproptosis-resistance program. Neutralizing co-induced IL6 further enhances PD-1 blockade efficacy. The treated TME transcriptionally recapitulates the T cell and myeloid signatures of pembrolizumab-responsive human patients, positioning LNP-IFNα2 as an off-the-shelf nanomedicine for ICI-resistant CRC.

## RESULTS

### LNP-mIFNα2 transgene selectively transfects tumor cells to express IFNα2 locally to overcome colon tumor lung metastasis resistance to PD-1 blockade immunotherapy

Lipid nanoparticle-encapsulated mouse codon usage-optimized IFNα2-encoding nanoplasmid^51^ (LNP-mIFNα2) gene immunotherapy selectively delivers the nanoplasmid to tumor-bearing lungs in mice^50^. To determine the target cells of LNP-mIFNα2 in the metastases-bearing lungs, LNP-mIFNα2 was administered to CT26 tumor-bearing mice using an experimental lung metastasis model (Fig. 1A-C). Single cells were prepared from the tumor-bearing lungs and CD45^-^PD-L1^+^ tumor cells, T (CD3^+^), and myeloid (CD11b^+^) cells were sorted out (Fig. 1C). Codon-usage-optimized mIFNα2 sequence-specific PCR analysis^50^ detected transgene expression predominantly in tumor cells, with lower expression in T and myeloid cells (Fig. 1D-E).

**Figure 1.**
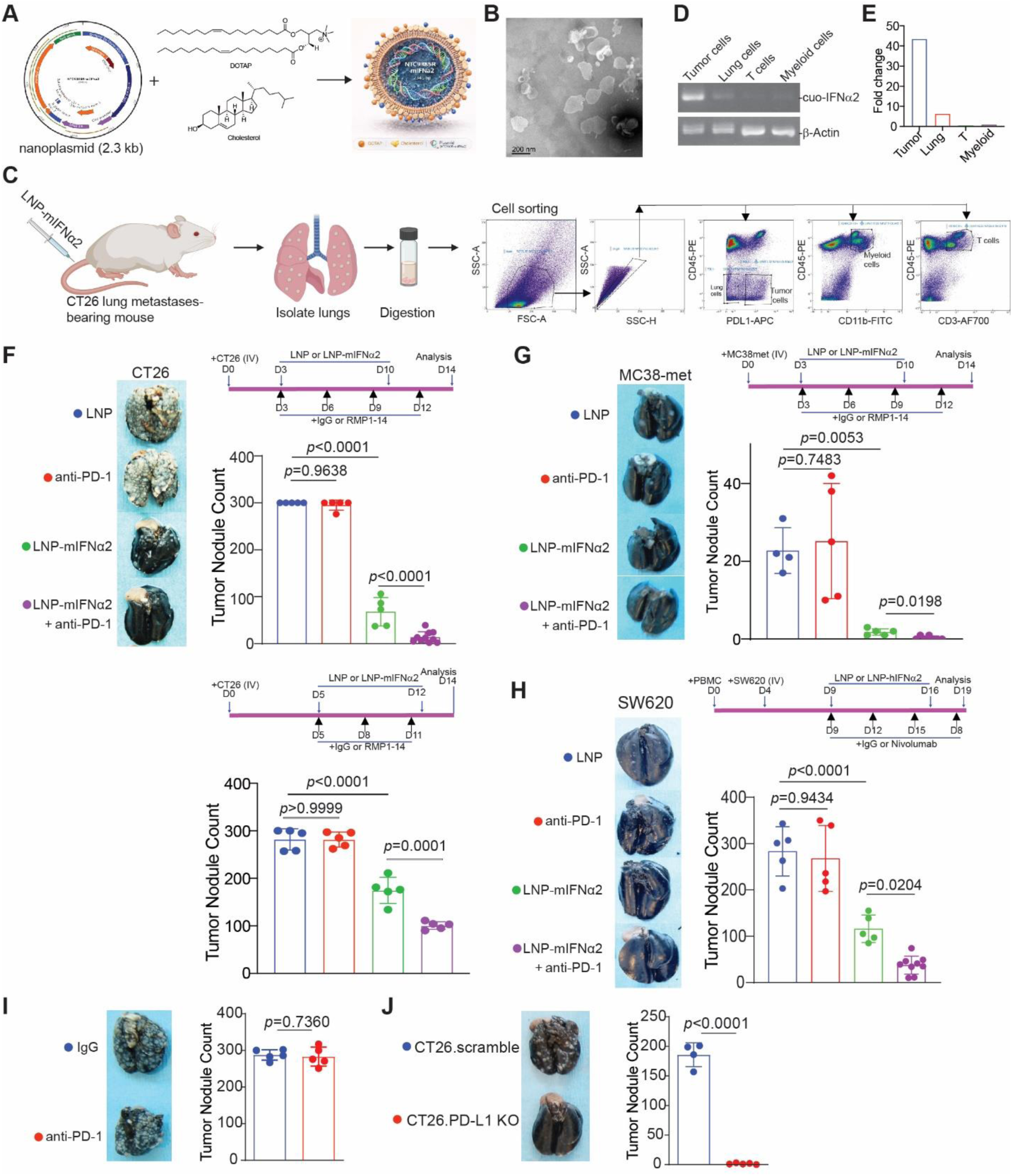
LNP-mIFNα2 gene immunotherapy forces tumor cells to express IFNα2 to overcome colon tumor resistance to PD-1 inhibitor immunotherapy. **A.** Schematic of LNP-mIFNα2 formulation. Codon-usage-optimized mouse IFNα2-encoding nanoplasmid NTC9385R was formulated with a DOTAP-cholesterol lipid to assemble lipid nanoparticles (LNP-mIFNα2). **B.** Scanning electron microscopy image of LNP-mIFNα2. Scale bar, 200 nm. **C.** Flow cytometry gating strategy for isolation of T (CD45^+^CD3^+^), myeloid (CD45^+^CD11b^+^) and tumor (CD45^-^PD-L1^+^) cells from CT26 metastases-bearing lungs. **D.** PCR detection of codon usage-optimized mIFNα2 transgene (cuo-IFNα2) in tumor cells, tumor-bearing total lung tissues (lung cells), T cells (CD3^+^), and myeloid cells (CD11b^+^) as shown in C. β-Actin serves as a loading control. **E.** Quantification of cuo-mIFNα2 transgene expression across sorted cell populations as shown in D. The expression level in myeloid cells was arbitrarily set at 1 and used as reference for other cell types. **F.** Antitumor efficacy of LNP-mIFNα2 alone and in combination with anti-PD-1 antibody in the CT26 syngeneic lung metastasis model. CT26 cells were injected intravenously into BALB/c mice. Mice were treated with LNP, anti-PD-1, LNP-mIFNα2, or LNP-mIFNα2 + anti-PD-1 beginning on day 3 (upper panel) or day 5 (lower panel) post-tumor inoculation. Upper panel, left: representative gross photographs of metastasis-bearing lungs at experimental endpoint of the day 3 treatment; top right: treatment scheme; bottom right: lung metastasis nodule counts for day 3 treatment initiation cohort. Lower panel, top: treatment scheme; bottom: nodule counts for day 5 treatment initiation cohort. Statistical comparisons in F-H by one-way ANOVA with Tukey’s post-hoc test; p-values shown. **G.** Antitumor efficacy in the MC38-met syngeneic lung metastasis model. MC38-met cells were injected intravenously into C57BL/6 mice. Mice were treated with LNP, anti-PD-1, LNP-mIFNα2, or LNP-mIFNα2+anti-PD-1 beginning on day 3 post-tumor inoculation as in F. Left: representative gross photographs of metastasis-bearing lungs. Right top: experiment scheme; right bottom: lung metastasis nodule counts. Data represent mean ± SD. **H.** Antitumor efficacy of LNP-hIFNα2 in a humanized NSG mouse model of human colon cancer lung metastasis. Tumor-bearing humanized mice were treated with LNP, nivolumab, LNP-hIFNα2 (LNP-encapsulated human IFNα2-encoding plasmid), or LNP-hIFNα2+nivolumab. Left: representative gross photographs of metastasis-bearing lungs. Right top: experiment scheme; right bottom: lung metastasis nodule counts. Data represent mean±SD. **I.** Mice were treated with anti-PD-1, CT26 cells were then injected intravenously into mice 1 hour later, followed by treatment with IgG or anti-PD-1 every 3 days for 4 more times. Left: representative gross photographs of metastases-bearing lungs at experimental endpoint. Right: lung metastases nodule counts. **J.** CT26.scramble and CT26.PD-L1 KO cells were injected to mice intravenously. Left: representative gross photographs of metastases-bearing lungs at experimental endpoint. Right: lung metastases nodule counts. Statistical comparison by two-tailed Student’s t-test; p-value shown. Data represent mean±SD.

We next evaluated antitumor efficacy in mouse colon tumor syngeneic lung metastasis models, which are resistant to PD-(L)1 ICI monotherapy^52,53^. Intravenous injection of CT26 cells followed by treatment with LNP-mIFNα2 alone or in combination with anti-PD-1 mAb reduced lung metastasis, while anti-PD-1 monotherapy had no effect (Fig. 1F). When treatment was delayed to day 5 to resemble a higher tumor burden, both LNP-mIFNα2 and the combination remained highly effective (Fig. 1F). To assess generalizability across colon tumor models, we tested LNP-mIFNα2 in the MC38-met model, a metastatic variant of the MC38 colon adenocarcinoma cell line^28^. Anti-PD-1 monotherapy was again ineffective, while LNP-mIFNα2 alone and in combination with anti-PD-1 significantly reduced lung metastasis (Fig. 1G). To evaluate translational potential, we established a humanized NSG mouse model by engrafting human peripheral blood mononuclear cells (PBMCs), followed by intravenous injection of the human colon cancer cell line SW620. Treatment with LNP-hIFNα2 alone and in combination with nivolumab significantly suppressed SW620 lung metastasis, while nivolumab alone had no significant effect (Fig. 1H). Two further observations define the resistance phenotype of this model. First, PD-1 blockade failed even when initiated before tumor cells seeded the lung. Mice were given anti-PD-1 antibody 1 h before intravenous CT26 injection and every 3 days thereafter; lung metastatic burden in these mice was indistinguishable from that of IgG-treated mice (Fig. 1I). Resistance in this model therefore does not arise from initiating checkpoint blockade too late relative to metastatic outgrowth. Second, the PD-(L)1 axis is nonetheless functionally important here: CT26 cells in which PD-L1 was deleted (CT26.PD-L1 KO) produced no lung metastases compared with CT26.scramble control cells (Fig. 1J). Tumor cell PD-L1 thus drives metastatic outgrowth, yet antibody blockade of PD-1 does not reproduce the effect of removing it. Together, these results establish that LNP-IFNα2 overcomes colon cancer resistance to PD-1 blockade immunotherapy to suppress colon cancer lung metastasis across multiple syngeneic and humanized preclinical models.

### LNP-mIFNα2 induces a transient cytokine response

Systemic IFNα2 protein production was detected in serum which peaked within 4-12 hours of LNP-mIFNα2 treatment and declined to near baseline by day 3 (Fig. 2A). In tumor-bearing lungs, mIFNα2 protein reached plateau at 12 hours and remained detectable through day 9 (Fig. 2B), demonstrating a durable local IFNα2 production within the metastatic site.

**Figure 2.**
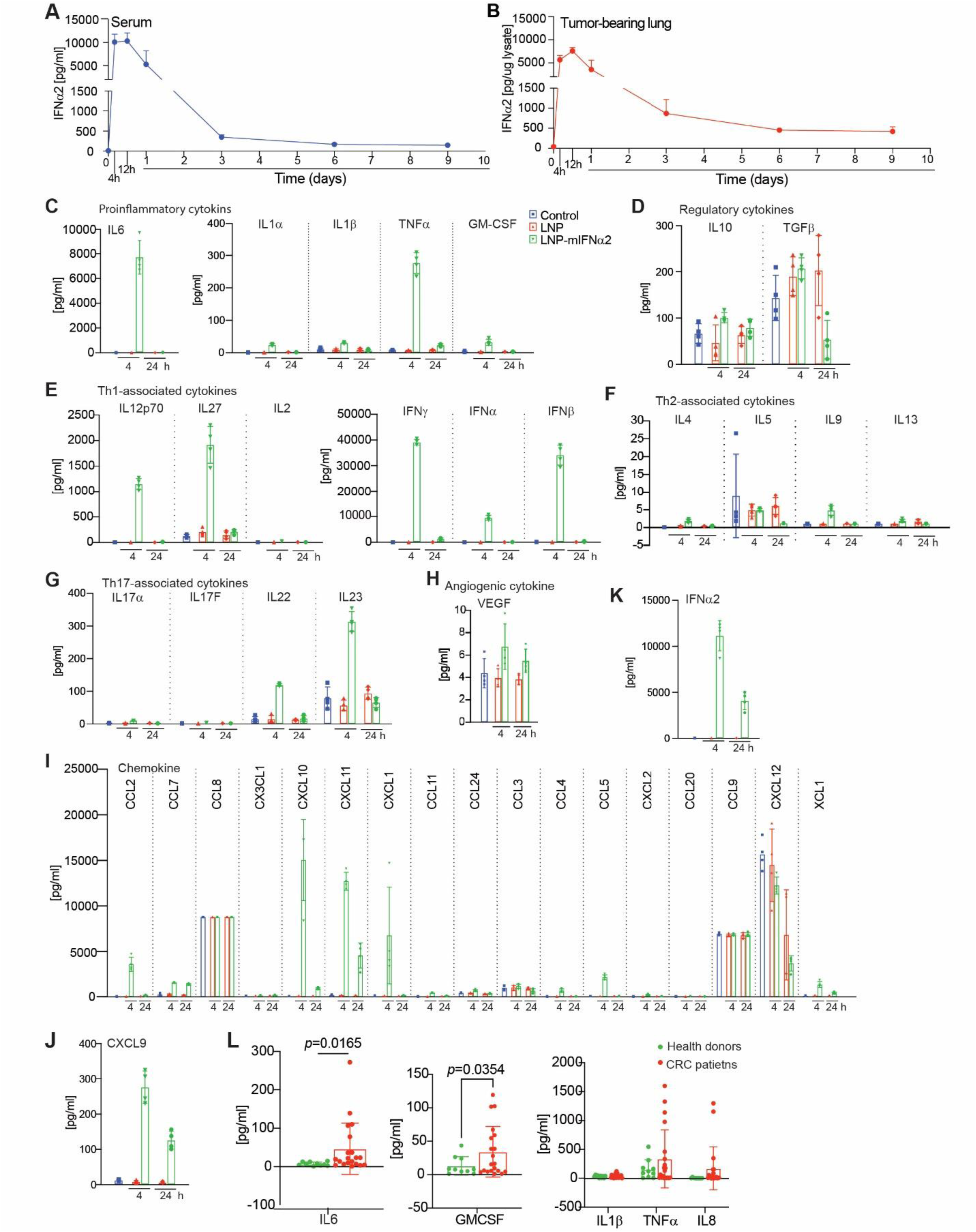
LNP-mIFNα2 induces a transient type I interferon and chemokine response. **A.** Serum mIFNα2 protein kinetics after intravenous LNP-mIFNα2 administration in CT26 metastasis-bearing mice, measured by ELISA. Data represent mean±SD. **B.** mIFNα2 protein kinetics in tumor-bearing lung tissue after intravenous LNP-mIFNα2 administration, measured by ELISA. Data represent mean±SD. **C.** Pro-inflammatory cytokines in mouse serum. Mice were untreated (Control) or treated with LNP or LNP-mIFNα2, and serum was collected 4 and 24 h after treatment. Each dot represents measurement from one mouse. Data represent mean±SD. **D-H**. Serum cytokines are measured by multiplex immunoassay as in C and grouped by functional class. **I.** Chemokines are measured by multiplex immunoassay as in C. **J.** Serum CXCL9 measured as in C. **K.** Serum IFNα2 protein levels at 4 h and 24 h after LNP or LNP-mIFNα2 administration measured by mouse IFNα2-specific ELISA. **L.** Serum specimens were collected from healthy donors (n=10) and colorectal cancer patients (n=20), and the indicated pro-inflammatory cytokines were measured by multiplex immunoassay.

A major adverse effect of lipid nanoparticle (LNP)-based nucleic acid therapeutics is the induction of systemic inflammatory cytokines following administration^54^. To characterize the immune response elicited by LNP-mIFNα2, serum was collected 4 and 24 h after treatment and analyzed using a multiplex cytokine assay. Among the pro-inflammatory cytokines, IL6 was markedly elevated at 4 h in LNP-mIFNα2-treated mice and declined toward baseline by 24 h (Fig. 2C), indicating an acute but self-limiting inflammatory response. In contrast, the classical inflammatory mediators IL1α, IL1β, TNFα, and GM-CSF remained at low levels, suggesting that LNP-mIFNα2 does not trigger a broad systemic inflammatory cytokine storm. Regulatory cytokines exhibited only modest changes following treatment (Fig. 2D). IL10 was transiently increased at 4 h and declined by 24 h, whereas TGFβ showed only modest fluctuations among treatment groups. Th1-associated cytokines displayed a selective activation profile (Fig. 2E). IL12p70 and IL27 were robustly induced at 4 h in LNP-mIFNα2-treated mice and decreased by 24 h, whereas IL2 remained unchanged. IFNα (IFNα1, IFNα6, and IFNα9) was dramatically elevated following LNP-mIFNα2 administration and returned to baseline by 24 h. IFNγ was also strongly induced at 4 h, consistent with rapid activation of NK cells and/or T cells downstream of type I IFN signaling. IFNβ showed a more modest increase. In contrast, Th2-associated cytokines (IL4, IL5, IL9, and IL13) remained at baseline across all treatment groups and time points (Fig. 2F), indicating that LNP-mIFNα2 preferentially activates type 1 rather than type 2 immune responses. Among Th17-associated cytokines, IL17A and IL17F were not induced, whereas IL22 was modestly increased and IL23 was markedly elevated at 4 h following LNP-mIFNα2 treatment before declining at 24 h (Fig. 2G). VEGF levels remained largely unchanged (Fig. 2H), indicating that systemic angiogenic cytokine production was not altered by treatment.

LNP-mIFNα2 also induced a distinct chemokine program (Fig. 2I). The IFN-inducible chemokines CXCL10 and CXCL11, together with CXCL1, CCL2, CCL7, CCL9, and CXCL2, were elevated, although with different kinetics. CXCL10, CXCL11, CCL9, and CCL2 were most prominent at 4 h, whereas CXCL1 remained elevated at 24 h. CXCL10, a canonical type I IFN-stimulated chemokine, promotes recruitment of CXCR3**⁺** effector T cells and NK cells, whereas CCL2 recruits inflammatory monocytes, supporting rapid innate immune activation following LNP-mIFNα2 administration. In contrast, CCL3, CCL4, CCL5, CCL11, CCL20, CCL24, CX3CL1, and CXCL12 were minimally affected. XCL1 was modestly increased, consistent with early activation of cytotoxic lymphocytes capable of recruiting XCR1⁺ cDC1s. CXCL9, an IFN-inducible chemokine that mediates T cell recruitment to lung metastases, was markedly elevated at 4 h and remained elevated at 24 h (Fig. 2J). Serum IFNα2 protein was at high levels at 4 h and remained detectable by 24 h (Fig. 2K), consistent with the pharmacokinetic profile observed in tumor-bearing lungs (Fig. 2A-B).

LNP-mIFNα2 boost therapy induces a similar cytokine profile as the prime therapy (Fig. S1A-I). Collectively, these findings demonstrate that LNP-mIFNα2 elicits a rapid but transient type I IFN-dominated cytokine and chemokine response characterized by induction of immune cell-recruiting chemokines without broad activation of pro-inflammatory cytokines.

To contextualize these mouse findings in human cancer, we analyzed serum cytokines and from CRC patients and healthy donors. CRC patients exhibited significantly elevated IL6 and GM-CSF (Fig. 2L), IL12p70 (Fig. S2A), IL10 (Fig. S2C), and the chemokines CXCL10 and CXCL5 (Fig. S2E), while Th2-and Th17-associated cytokines were unchanged (Fig. S2B & D). These findings are consistent with the phenomenon that human CRC is characterized by a persistent, often low-grade inflammatory tumor microenvironment^55^.

### LNP-mIFNα2 induces a transient and self-resolving intrapulmonary neutrophil response

To assess the pathological changes in the lungs, H&E-stained lung sections were evaluated by board-certified pathologists. In untreated control mice, lung sections showed preserved alveolar architecture with mild emphysematous changes and evidence of poorly differentiated parenchymal malignancy consistent with CT26 metastatic disease (Fig. S3A). In LNP-treated mice, the histological appearance at 4 h was largely unchanged, sections showed benign lung tissue or mild emphysematous changes (Fig. S3B). By 24 h after LNP treatment, mild to moderate emphysematous changes and focal perivascular lymphocytic infiltration were noted, without significant neutrophilia (Fig. S3C), consistent with very mild, resolving vehicle-related changes.

LNP-mIFNα2-treated mice displayed a distinct, time-dependent neutrophil response. At 4 h, average intra-alveolar neutrophil counts were elevated approximately twice the control baseline and 2.6-fold above the LNP 4 h level, indicating a robust but compartmentalized neutrophil influx into the alveolar space (Fig. S3B, S3E). Higher-magnification images of LNP-mIFNα2-treated lungs at 4 h confirmed the presence of neutrophils, characterized by multi-lobed nuclei, accumulating within alveolar spaces (Fig. S3D), adjacent to structurally intact bronchioles and pulmonary arteries, without evidence of vessel wall disruption, necrosis, diffuse alveolar damage, or parenchymal tissue injury. Despite the elevated neutrophil count, pathological evaluation noted no evidence of tissue injury or inflammation at this timepoint. By 24 h post-LNP-mIFNα2 injection, alveolar neutrophil counts resolved substantially to a level that is comparable to the untreated control level, demonstrating rapid clearance of the early neutrophil influx (Fig. S3C & D, S3E). Histologically, LNP-mIFNα2-treated lungs at 24 h showed only mild emphysematous changes in some sections, with benign lung tissue in others, and no evidence of persistent tissue injury or inflammation (Fig. S3C & D). LNP-mIFNα2 boost therapy induced a similar pathological change as the prime treatment in tumor-bearing mouse lungs (Fig. S4A-D).

To validate the histological neutrophil influx, flow cytometry was performed on tumor-bearing lungs after prime and boost treatment. LNP-mIFNα2 increased neutrophil frequencies at 4 h post-prime and post-boost, returning toward baseline by 24 h (Fig. 3A-H). This complementary cell-based quantification confirms the transient, self-resolving neutrophil response observed by H&E histopathology.

**Figure 3.**
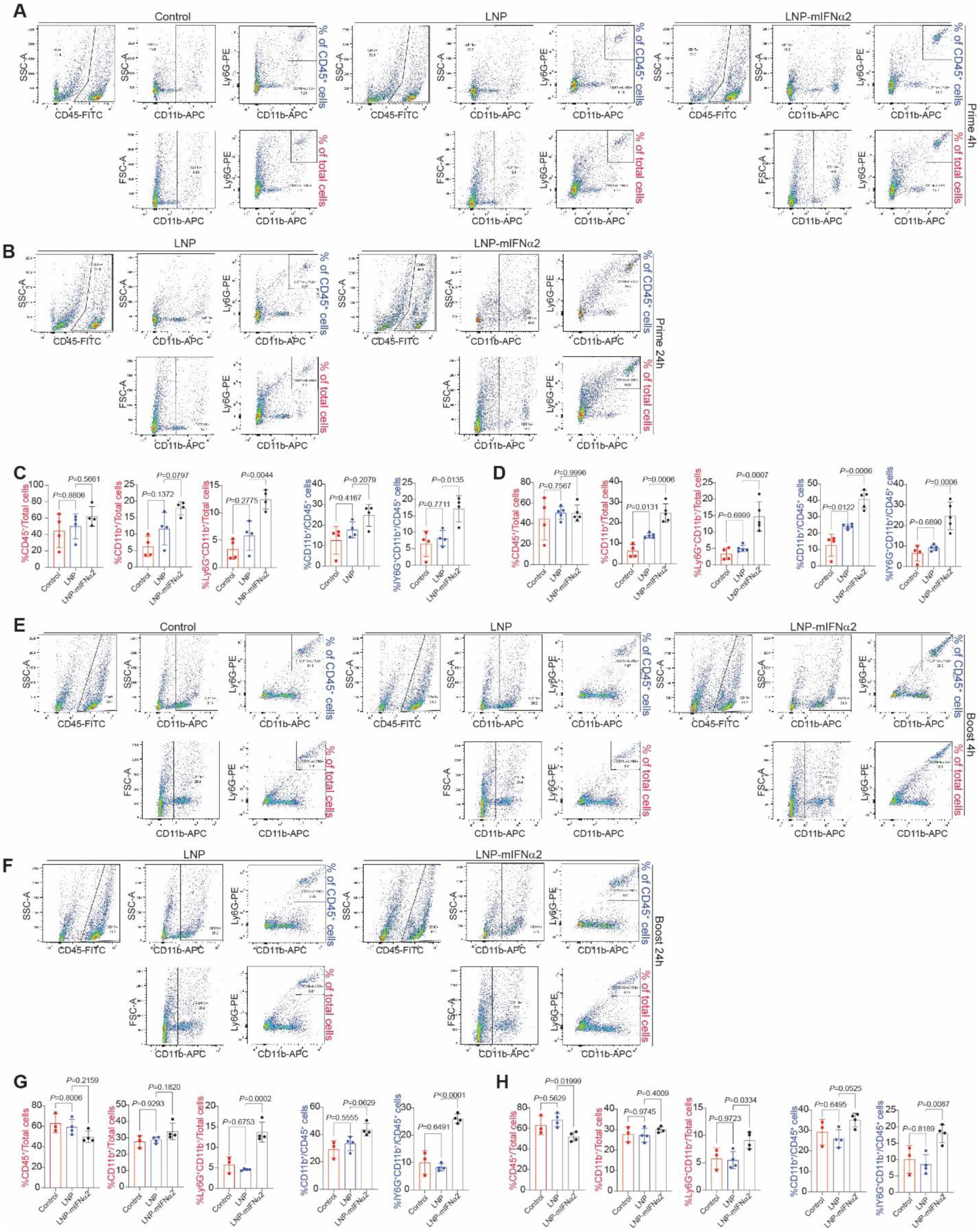
LNP-mIFNα2 treatment expands neutrophils in metastases-bearing lungs in mice. A-B. Representative flow cytometry gating strategy for tumor-bearing lung leukocytes after prime treatment. Tumor-bearing mice were either untreated (Control), treated with LNP or LNP-mIFNα2. Lungs were collected at 4h (A) and 24h (B) later and analyzed by flow cytometry. **C-D.** Quantification of the indicated cell types at 4 (C) and 24 (D) hours after the prime treatment as shown in A and B. Each dot represents one mouse. Data represent mean±SD. **E-F.** Representative flow cytometry gating strategy after boost treatment. Lung metastases-bearing mice as shown in A-B were treated again (boost) 7 days later. Lungs were collected 4 (E) and 24 (F) hours later and analyzed by flow cytometry. **G-H.** Quantification of the indicated cell types at 4 (G) and 24 (H) hours after boost.

Collectively, the pathological and flow cytometric analyses demonstrate that LNP-mIFNα2 immunotherapy induces a transient alveolar neutrophil influx that peaks at 4 h and resolves by 24h after both prime and boost dosing, without tissue injury, diffuse alveolar damage, vascular disruption, or persistent pulmonary inflammation (Figs. 3, S3, S4). The convergence of histological and flow cytometric findings across two complementary approaches confirms that this neutrophil response is robust, reproducible, and self-limiting. This profile is consistent with the serum cytokine data (Figs. 2 & S1) showing a self-resolving innate immune response and supports the absence of overt acute pulmonary injury under the conditions tested.

**Figure 4.**
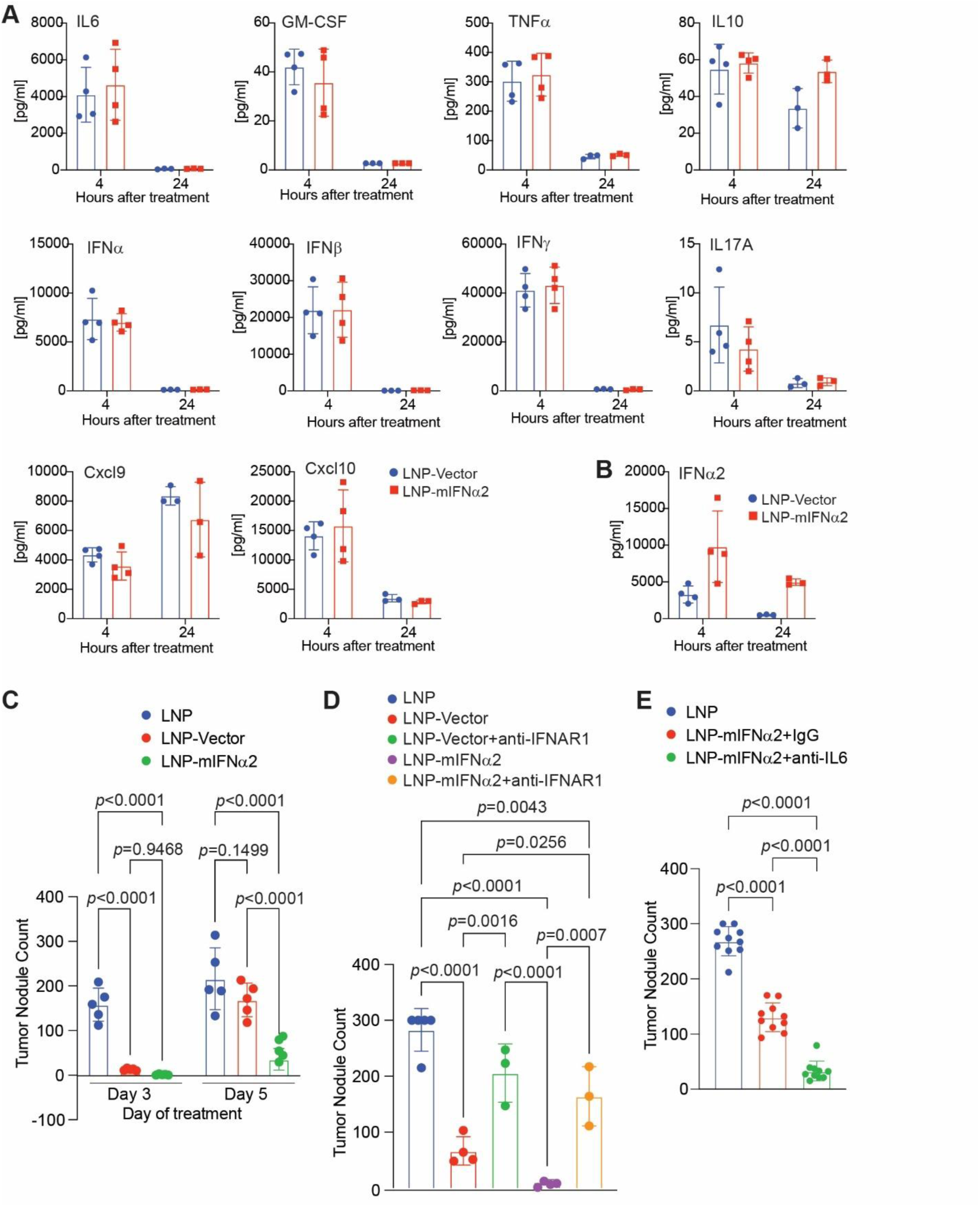
LNP-mIFNα2 suppresses CT26 pulmonary metastases through IFNAR1 signaling and is associated with IL6 induction. **A.** Serum cytokine profiles in CT26 metastases-bearing mice. Mice were treated intravenously with lipid nanoparticle (LNP)-encapsulated empty vector (LNP-Vector) or LNP-mIFNα2. Serum concentrations of IL6, GM-CSF, TNFα, IL-10, IFNα, IFNβ, IFNγ, IL-17A, Cxcl9, and Cxcl10 were measured at 4 h and 24 h after treatment. Data are shown as mean±SD. **B.** Serum IFNα2 protein levels in mice treated as in (A), measured at 4 h and 24 h after treatment. Data are shown as mean±SD. **C.** Quantification of pulmonary metastatic nodule number in CT26 metastases-bearing mice treated with LNP, LNP-Vector, or LNP-mIFNα2 administered at Day 3 or Day 5 after tumor cell injection. Each dot represents one mouse. Data are shown as mean±SD. **D.** Quantification of pulmonary metastatic nodule number in CT26 metastases-bearing mice treated with LNP-Vector or LNP-mIFNα2 in the presence of IgG or anti-IFNAR1 monoclonal antibody. Each dot represents one mouse. Data are shown as mean±SD. **E.** Quantification of pulmonary metastatic nodule number in CT26 metastases-bearing mice treated with LNP or LNP-mIFNα2. The LNP-mIFNα2-treated group was further divided and treated with IgG or anti-IL6 neutralizing monoclonal antibody administered once prior to LNP-mIFNα2 treatment. Each dot represents one mouse. Data are shown as mean±SD.

### LNP-mIFNα2 induces antitumor efficacy through IFNAR1 signaling

Systemic administration of LNP-mRNA elicits a viraemia-like surge in IFN-I and innate immune activation^19^. However, LNP-mediated delivery of plasmid DNA (pDNA) introduces an additional layer of innate immune stimulation, driving rapid production of IFN-I and pro-inflammatory cytokine IL6^54^. We therefore hypothesized that LNP-mIFNα2, which delivers a pDNA transgene encoding IFNα2, would engage both the innate response inherent to pDNA delivery and the direct IFNAR1-mediated antitumor activity of the encoded IFNα2 protein, and that this dual mechanism of IFN-I induction would confer superior antimetastatic efficacy. At 4 h after injection, LNP-mIFNα2-treated mice showed elevated serum levels of IL6, GM-CSF, TNFα, IFNα, IFNβ, IFNγ, Cxcl9, CXCL10, IL10, and IL17A relative to LNP-treated mice, with most cytokines declining substantially by 24 h. Notably, LNP-Vector-treated mice displayed a comparable cytokine induction at 4 h (Fig. 4A). This inflammatory response elicited by LNP-pDNA is consistent with cGAS-STING-mediated sensing of the encapsulated plasmid DNA backbone, which drives IFN-I production independently of transgene expression^54,56^. In line with active IFNα2 transgene expression, serum IFNα2 protein was significantly elevated only in LNP-mIFNα2-treated mice at 4 h and remained detectable at 24 h, whereas LNP-Vector-treated animals showed no measurable IFNα2 protein (Fig. 4B), confirming that LNP-mIFNα2 drives a quantitatively and qualitatively distinct IFN response beyond baseline pDNA-mediated innate activation.

To assess whether the augmented IFN response of LNP-mIFNα2 translates into antitumor efficacy, we quantified pulmonary metastatic burden following treatment at Day 3 or Day 5 post-tumor injection, representing early and late metastatic stages, respectively (Fig. 4C). At Day 3, both LNP-Vector and LNP-mIFNα2 significantly reduced lung metastatic nodule counts relative to LNP control and did not differ from each other, indicating that the innate response to the plasmid backbone alone is sufficient to control early, low-burden metastases. At Day 5, in contrast, LNP-Vector no longer significantly reduced metastatic burden relative to LNP control, whereas LNP-mIFNα2 remained highly effective and was significantly superior to LNP-Vector. The therapeutic advantage of LNP-mIFNα2 over LNP-Vector therefore emerges specifically at Day 5, a time point that mimics high tumor burden in the clinical setting, suggesting that robust transgene-driven IFNα2 expression and direct IFNAR1 signaling are especially critical when the immunosuppressive tumor microenvironment is more established, consistent with observations that mRNA-LNP-induced IFN-I is required to sensitize immunologically cold tumors to checkpoint blockade^19^. To confirm IFNAR1 dependence, mice were treated with anti-IFNAR1 blocking antibody during treatment. IFNAR1 blockade diminished the antitumor effect of LNP-Vector and LNP-mIFNα2 (Fig. 4D). These results demonstrate that the antitumor efficacy of LNP-Vector and LNP-mIFNα2 is dependent on canonical IFNAR1 signaling and is quantitatively superior to the cGAS-STING-mediated baseline inflammatory response to the vector backbone, particularly under conditions of high tumor burden.

### Neutralization of IL6 enhances LNP-mIFNα2 antitumor therapeutic activity

Among the cytokines transiently induced by LNP-mIFNα2, IL6 was among the most significantly elevated at 4 h and rapidly resolved by 24 h (Figs. 2C & 4A). This acute IL6 surge is characteristic of LNP-pDNA-induced inflammation driven by cGAS-STING activation, which prominently upregulates IL6^54^. To determine the functional consequence of this transient IL6 response on antitumor activity, we treated CT26 metastases-bearing mice with LNP-mIFNα2 in combination with either isotype control (IgG) or an anti-IL6 neutralizing monoclonal antibody administered once prior to LNP-mIFNα2 (Fig. 4E). Neutralization of IL6 significantly enhanced the antitumor efficacy of LNP-mIFNα2, resulting in a further reduction of pulmonary metastatic nodule counts compared with LNP-mIFNα2+IgG controls. These observations indicate that the transient IL6 surge induced by LNP-mIFNα2 is not required for its antitumor effect and may instead dampen therapeutic activity, a known downstream consequence of acute IL6 elevation in the tumor microenvironment^57^. Blocking this IL6 burst therefore unmasks an augmented antitumor response, positioning transient IL6 as a modulatable liability of pDNA-LNP-based IFNα2 therapy. Together, these findings suggest that co-administration of IL6 blockade with LNP-mIFNα2 represents a rational combinatorial strategy to maximize the antimetastatic efficacy of IFN-I-based nanoparticle targeted immunotherapy.

### LNP-mIFNα2 broadly reprograms the tumor-infiltrating immune compartments toward an antitumor immune phenotype

To probe the effect of LNP-mIFNα2 on the tumor microenvironment (TME), we isolated CD45^+^ cells from CT26 tumor lung metastases (Fig. S5A & B) and performed single cell RNA sequencing (scRNA-seq). After integration and clustering, cells were annotated into major immune and residual stromal populations based on differentially expressed genes and canonical marker genes (Fig. S5C-F). Inspection of the split UMAP and proportion plot revealed striking differences in both the proportion of monocytes versus macrophages and in the composition of the T cell compartment between conditions (Fig. 5A). In the LNP control tumor, there were high Tumor-associated Macrophages (TAMs) and low Ly6^+^ inflammatory monocytes. In contrast, the LNP-mIFNα2-treated tumor harbored low TAMs and high monocytes (Fig. 5A). We also observed an expansion of Tpex-like T cells in the LNP-mIFNα2-treated tumor (Fig. 5A). To investigate these differences at higher resolution, we separately subclustered the monocyte/macrophage and T cell compartments.

**Figure 5.**
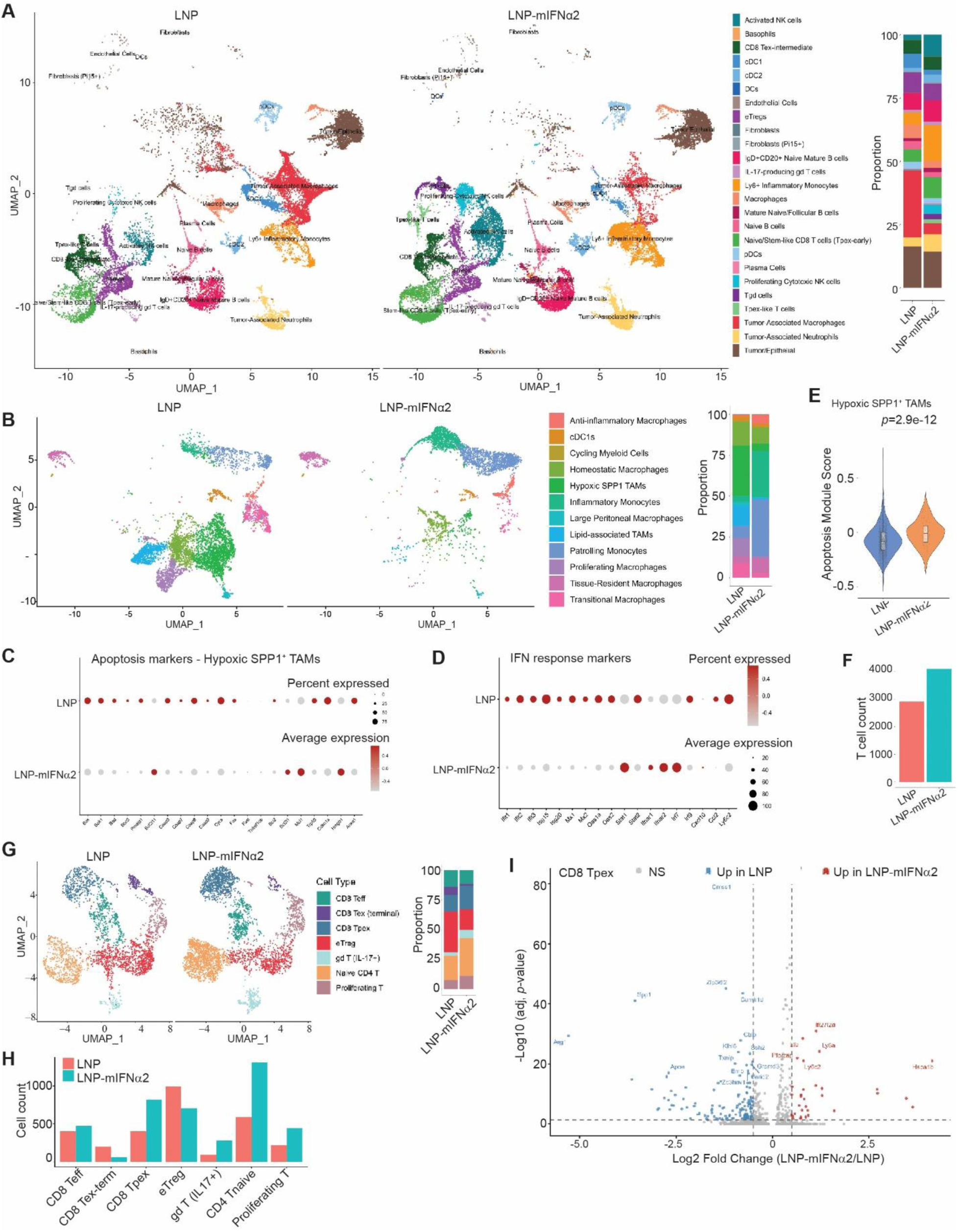
LNP-mIFNα2 reshapes the tumor immune microenvironment in lung metastases. **A.** UMAP visualization of integrated single-cell RNA sequencing data showing CD45⁺ tumor-infiltrating immune cells from lung metastases in mice treated with LNP control or LNP-mIFNα2. Stacked bar graph (right) shows the relative abundance of each immune cell population in the two treatment groups. **B.** UMAP of re-clustered monocyte/macrophage populations (derived from the original Ly6⁺ inflammatory monocyte, macrophage, and tumor-associated macrophage clusters). Stacked bar graph (right) shows the relative abundance of each myeloid subtype in LNP control and LNP-mIFNα2-treated samples. **C.** Dot plot of apoptosis-associated gene expression in Hypoxic SPP1^+^ TAMs, split by sample (LNP vs. LNP-mIFNα2). **D.** Dot plot of IFN-response marker gene expression in inflammatory monocytes and patrolling monocytes clusters. **E.** Violin plot showing apoptosis module scores in hypoxic SPP1⁺ tumor-associated macrophages from LNP control and LNP-mIFNα2-treated tumors. Statistical significance was determined using the Wilcoxon rank-sum test (pseudoreplication). **F.** Bar graph showing the total number of T cells recovered from each scRNA-seq sample. **G.** UMAP of re-clustered T-cell populations (derived from the original CD8 Tex-intermediate, eTreg, IL-17-producing γδ T cell, naïve/stem-like CD8 T cell [Tpex-early], Tgd, and Tpex-like T-cell clusters). Stacked bar graph (right) shows the relative abundance of each T-cell subtype in LNP control and LNP-mIFNα2-treated samples. **H.** Bar graph showing the absolute number of cells in each re-clustered T-cell subtype in LNP control and LNP-mIFNα2-treated tumors. **I.** Volcano plot showing differentially expressed genes in the CD8 Tpex cell population comparing LNP-mIFNα2-treated and LNP control tumors. Red, genes upregulated in LNP-mIFNα2; blue, genes upregulated in LNP control.

### LNP-mIFNα2 reprogrammed the monocyte/macrophage compartment in lung metastasis

The three monocyte/macrophage clusters from the overall UMAP (Ly6^+^ Inflammatory Monocytes, Macrophages, and Tumor-Associated Macrophages) were subsetted and re-clustered. Non-myeloid clusters identified by expression of lineage-specific markers were excluded prior to downstream analysis (Fig. S6A, B). The remaining clusters were annotated into 12 cell types using a combination of differential gene expression, canonical marker genes, and pairwise DEG analysis between selected clusters (Fig. S6B).

Quantification of myeloid subtype proportions revealed that the broad shift from macrophages to monocytes observed in the overall UMAP (Fig. 5A) was attributable specifically to a reduction in Hypoxic SPP1 TAMs, a population of myeloid cell with potent immune suppressive function ^58–61^, and Lipid-associated TAMs, with a corresponding increase in Inflammatory and Patrolling Monocytes in the LNP-mIFNα2 group (Fig. 5B). In the LNP control, Patrolling Monocytes, Inflammatory Monocytes, and Hypoxic SPP1^+^ TAMs all expressed higher levels of Arginase 1 (*Arg1*), a key immunosuppressive enzyme in the TME. Hypoxic SPP1 TAMs additionally expressed higher levels of *Spp1* in LNP control tumors (Fig. S6C). We have previously shown that IRF8 represses *Spp1* expression in myeloid cells^28^. Given that LNP-mIFNα2 elevates type I IFN levels in the TME, IRF8-mediated transcriptional repression of *Spp1* in myeloid cells may represent a key mechanism of LNP-mIFNα2 efficacy. Consistent with this, *Spp1* expression was globally repressed across myeloid cells in the LNP-mIFNα2 group (Fig. S6C). Hypoxic SPP1^+^ TAMs in the LNP-mIFNα2 group additionally downregulated suppressive markers including *Trem2* and *Arg1*, and upregulated IFN-response genes to among the highest levels of any myeloid subtype (Fig. S6C-E).

The striking reduction in Hypoxic SPP1^+^ TAMs in LNP-mIFNα2-treated tumors prompted us to investigate the fate of these cells. We hypothesized that SPP1^+^ TAMs in LNP-mIFNα2-treated tumors were undergoing apoptosis. We compared apoptotic gene expression in SPP1^+^ TAMs between conditions (Fig. 5C). SPP1^+^ TAMs exhibited meaningfully higher apoptotic module scores in LNP-mIFNα2-treated tumors compared to LNP controls (Fig. 5E), suggesting that LNP-mIFNα2 drives both transcriptional programming and apoptotic elimination of this immunosuppressive population.

In the Inflammatory Monocytes cluster, distinguished from Patrolling Monocytes by expression of *Sell*, *Itga4*, *Prtn3*, *Mmp8*, and *Fn1*, while Patrolling Monocytes expressed *Itgax*, *Cd300e*, *Lilra5*, *Cd36*, and *Vegfc* (Fig. S6F), LNP-mIFNα2 induced upregulation of interferon-inducible GTPase *Ligp1* and granzyme *Gzma* (Fig. S6E). Further analysis confirmed high expression of canonical IFN-response genes in the Inflammatory Monocytes of LNP-mIFNα2-treated tumors (Fig. 5D). Co-expression of *Prtn3*, *Elane*, and *Mmp8* in this population is consistent with emergency myelopoiesis generating granulocyte-monocyte precursor-derived monocytes that are subsequently reinforced toward an IFN-responsive identity by the LNP-mIFNα2 treatment. Direct comparison of the two monocyte clusters confirmed their distinct identities, with Inflammatory Monocytes enriched for granule and emergency myelopoiesis transcripts and Patrolling Monocytes enriched for patrolling/scavenger markers (Fig. S6F).

### LNP-mIFNα2 expanded the CD8 Tpex compartment and relieved Tpex quiescence

The elimination of SPP1^+^ TAMs and emergence of IFN-responsive monocytes, which are known to stimulate T cells in tumors^62,63^, in LNP-mIFNα2-treated tumors prompted us to examine whether myeloid reprogramming was accompanied by favorable changes in the T cell compartment. The six T cell clusters from the overall UMAP were subsetted and re-clustered using the same pipeline as for the myeloid compartment. Non-T cell clusters were identified and excluded (Fig. S7A, B) and the remaining cells were integrated using Harmony prior to final cluster annotation (Fig. S7C-F). LNP-mIFNα2 expanded the total T cell compartment (Fig. 5F).

LNP-mIFNα2 induced significant changes in T cell subtype composition (Fig. 5G & H). Terminally exhausted CD8 T cells (CD8 Tex-term) were reduced in the treatment group, while progenitor-exhausted CD8 T cells (CD8 Tpex), the subtype known to expand and respond to anti-PD-1 immunotherapy, increased approximately two-fold in number (Fig. 5H). Effector Tregs (eTregs) were reduced in LNP-mIFNα2, and Naïve CD4 T cells, γδ T cells, and Proliferating T cells were expanded (Fig. 5G & H). DEG analysis of the remaining T cell subtypes (CD8 Tex-interm, Tpex-early, eTregs, and IL17^+^ γδ T cells) similarly revealed upregulation of IFN-stimulated genes in the LNP-mIFNα2 group (Fig. S7G). DEG analysis of the CD8 Tpex cluster revealed that in LNP control tumors, CD8 Tpex are maintained in a quiescent state, characterized by elevated expression of the mRNA destablizer *Zfp36l2*, the metabolic checkpoint *Txnip*, and the immunosuppressive enzyme *Arg1* (Fig. 5I). LNP-mIFNα2 relieved this quiescent state and directed CD8 Tpex toward an IFN-response phenotype, evidenced by significant upregulation of IFN-stimulated genes *Ifi27l2a* and *Ly6a* (Fig. 5I).

### LNP-mIFNα2 expands cytotoxic NK cell subsets and shifts the B cell compartment toward plasma cell and mature states

In addition to the myeloid and T cell compartments, LNP-mIFNα2 reprogrammed the intratumoral B and NK cell populations. B cells showed proportional shifts toward Plasma Cells and mature B cell states in the LNP-mIFNα2 group, encompassing the major B cell subsets, including IgD^+^CD20^+^ naive mature B cells, naive B cells, plasma cells, and mature naive B cells (Fig. 5A). Activated NK cells and Proliferating Cytotoxic NK cells, identified from the overall UMAP, were both substantially expanded in proportion and in absolute number in the LNP-mIFNα2 group relative to LNP control (Fig. S8A-C). Differential gene expression analysis of these two NK subsets revealed broad upregulation of IFN-stimulated and cytotoxic effector transcripts in LNP-mIFNα2-treated tumors (Fig. S8D), consistent with the cytolytic reprogramming observed across other immune compartments. Together with the myeloid and T cell findings, these data indicate that LNP-mIFNα2 broadly reprograms the tumor immune microenvironment across innate and adaptive lymphoid compartments to promote antitumor immunity.

### LNP-mIFNα2 depletes high cycling tumor cells and reverses cuproptosis-resistance programs in tumor cells

To characterize the direct effects of LNP-mIFNα2 on tumor and stromal cell populations in the CT26 lung metastatic niche, we performed scRNA-seq on the CD45^-^ cells from LNP-and LNP-mIFNα2-treated mice. Because the LNP-mIFNα2-treated samples had reduced tumor cell content and enriched immune cells, CD45^+^ cells were removed by CD45 thresholding prior to analysis (LNP: 20,947 cells; LNP-mIFNα2: 7,097 cells after filtering, Fig. S9A & B). Supervised clustering resolved seven transcriptionally distinct populations, comprising five cancer cell states, Fibroblasts, and Endothelial cells (Fig. 6A & B, Fig. S9C & D), which were annotated based on mouse colon tumor marker genes^64^. Endothelial cells were excluded from the subsequent tumor cell subtype analyses. UMAP visualization and proportional composition analysis revealed striking LNP-mIFNα2-driven remodeling of the tumor cell landscape: the proportion of High Cycling CC was markedly reduced in LNP-mIFNα2-treated tumors, while the proportion of IFN Response CC was substantially increased (Fig. 6A-B).

**Figure 6.**
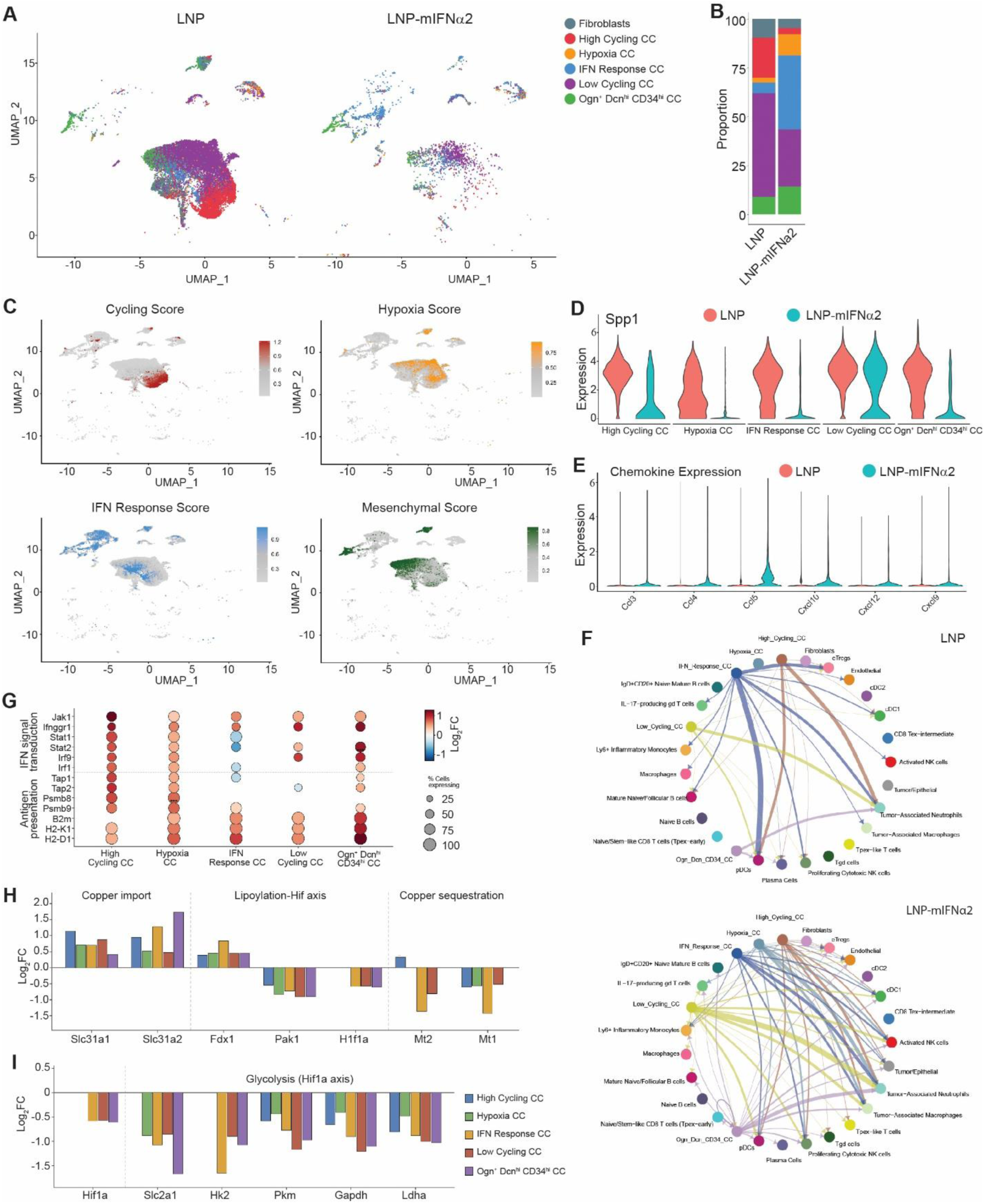
mIFNα2 transgene therapy drives depletion of high cycling cancer cell and reverses cuproptosis-resistance programs in tumor cells. **A.** UMAP of all CD45^−^ cells colored by annotated cluster identity split by sample. **B.** Stacked proportion bar plot of tumor cell populations as in (A). **C.** Feature plots of module scores for proliferation, hypoxia, IFN response, and mesenchymal gene programs. **D.** Violin plots split by sample of *Spp1* expression across tumor cell subtypes. **E.** Violin plots split by sample of chemokine gene expression across tumor cell subtypes. **F.** CellChat circle plots of chemokine network specifically/only showing tumor cells as the senders in LNP control (left) and LNP-mIFNα2 (right) **G.** DotPlot of IFN signal transduction genes and MHC I antigen-presentation genes across tumor cell subtypes, comparing LNP-mIFNα2 to LNP. **H.** Bar plot of average log2 fold-change (LNP-mIFNα2 vs LNP) for core cuproptosis pathway genes across the same tumor subtypes, grouped by function: copper import, the lipoylation/Hif axis, and copper sequestration. **I.** Bar plot of average log2 fold-change (LNP-mIFNα2 vs LNP) for genes of the HIF-1α-driven glycolytic (cuproptosis-resistance) signature in the indicated tumor cell subtypes.

Module scoring confirmed the compositional shifts observed by UMAP: the high cycling score was highest in High Cycling CC and was broadly reduced in LNP-mIFNα2, while the IFN response score was observed more broadly across cell populations (Fig. 6C). The expression of *SPP1*, an oncogenic driver in cancer cells^65,66^, was decreased across all tumor subtypes in LNP-mIFNα2-treated mice (Fig. 6D). In addition, tumor cells in LNP-mIFNα2-treated mice showed remodeled chemokine expression consistent with enhanced antitumor immune recruitment. *Ccl3*, *Ccl4*, *Ccl5*, *Cxcl9*, *Cxcl10*, and *Cxcl12*, which collectively recruit NK cells, dendritic cells, and CD8^+^ T cells, were elevated in the treatment group (Fig. 6E, Fig. S9E). To interrogate the functional consequence of chemokine expression remodeling, we performed receptor-ligand interaction analysis using CellChat, restricting the sender role to tumor cell subtypes. The tumor-cell derived chemokine network was substantially expanded in LNP-mIFNα2-treated mice relative to LNP controls, with all tumor cell subtypes showing increased number and strength of interactions with immune cell populations (Fig. 6F).

Given that the IFN signaling axis has recently been shown to directly transactivate *FDX1* and sensitize tumor cells to cuproptosis and overcome tumor cell resistance to cuproptosis^67–70^, we asked whether LNP-mIFNα2 engages this pathway in tumor cells. Across tumor cell subtypes, LNP-mIFNα2 treatment increased expression of IFN signal transduction components (*Jak1*, *Stat1*, *Stat2*, *Irf1*, *Irf9*, and the IFN-γ receptor *Ifngr1*) and the IRF1-dependent MHC class I antigen-presentation program (*Tap1/2*, *Psmb8/9*, *B2m*, *H2-K1/D1*) (Fig. 6G). This was accompanied by a consistent increase in *Fdx1*, the rate-limiting cuproptosis effector and direct IRF1 transcriptional target, across all tumor cell subtypes (Fig. 6H). A reciprocal and broad decrease in the copper-sequestering metallothionein *Mt1* (Fig. 6H) and downregulation of the Hif1a-regulated cuproptosis-resistance signature (Fig. 6I) was also observed in LNP-mIFNα2-treated tumors. This Fdx1-up/Mt1-down configuration is the converse of the signature associated with cuproptosis resistance in IFN-experienced tumors, in which BACH1 loss de-represses MT1E/MT1X to blunt cuproptosis despite active STAT1-IRF1 signaling^67,69,70^. Together with our finding that LNP-mIFNα2 increases copper-importer expression (*Slc31a1*, *Slc31a2*) across tumor subtypes (Fig. 6H), these data suggest that LNP-mIFNα2-induced IFN signaling sensitizes tumor cells to cuproptosis, raising the possibility that cuproptosis represents a previously unappreciated layer of tumor-intrinsic reprogramming by IFNα2 therapy.

### The immune landscape of LNP-mIFNα2-treated mouse tumors transcriptionally recapitulates the T cell and myeloid expansion signatures of PD-1 blockade responding human cancer patients

To determine whether LNP-mIFNα2-induced immune reprogramming in mouse tumors recapitulates human immunotherapy response programs, we analyzed a publicly available scRNA-seq dataset from breast cancer patients treated with neoadjuvant pembrolizumab^71^. Patients were stratified into T cell clonotype expanders (E), those exhibiting clonal T cell expansion upon pembrolizumab treatment, and non-expanders (NE), providing a human benchmark for immunotherapy-responsive versus-resistant tumor immune microenvironments. Analysis of the human T cell compartment resolved nine distinct phenotypes by UMAP, including Tpex, Terminal Exhausted CD8^+^, Proliferating T cells, and Tregs, which were enriched in E patients (Fig. 7A). Conversely, Naive/central memory CD4, Cytotoxic Effector CD8^+^, and Effector Memory CD8^+^ populations predominated in NE patients. Boxplot quantification confirmed significant enrichment of Tpex, Proliferating T cells, Tregs, and Terminal Exhausted CD8^+^ cells in E patients, whereas Naive/central memory CD4, Cytotoxic Effector CD8^+^, and Effector Memory CD8^+^ subsets were enriched in NE patients (Fig. 7B). Volcano plot analysis of the Tpex population revealed upregulation of effector and cytotoxic genes, including *GZMB*, *NKG7*, *CCL3*, *CCL4*, and *HOPX*, in E patients, closely mirroring the gene programs upregulated in LNP-mIFNα2-treated mouse CD8 T cells, where *Gzmb*, *Nkg7*, *Prf1*, and *Ccl5* were induced and Tpex-early populations expanded markedly (Fig. 7B).

**Figure 7.**
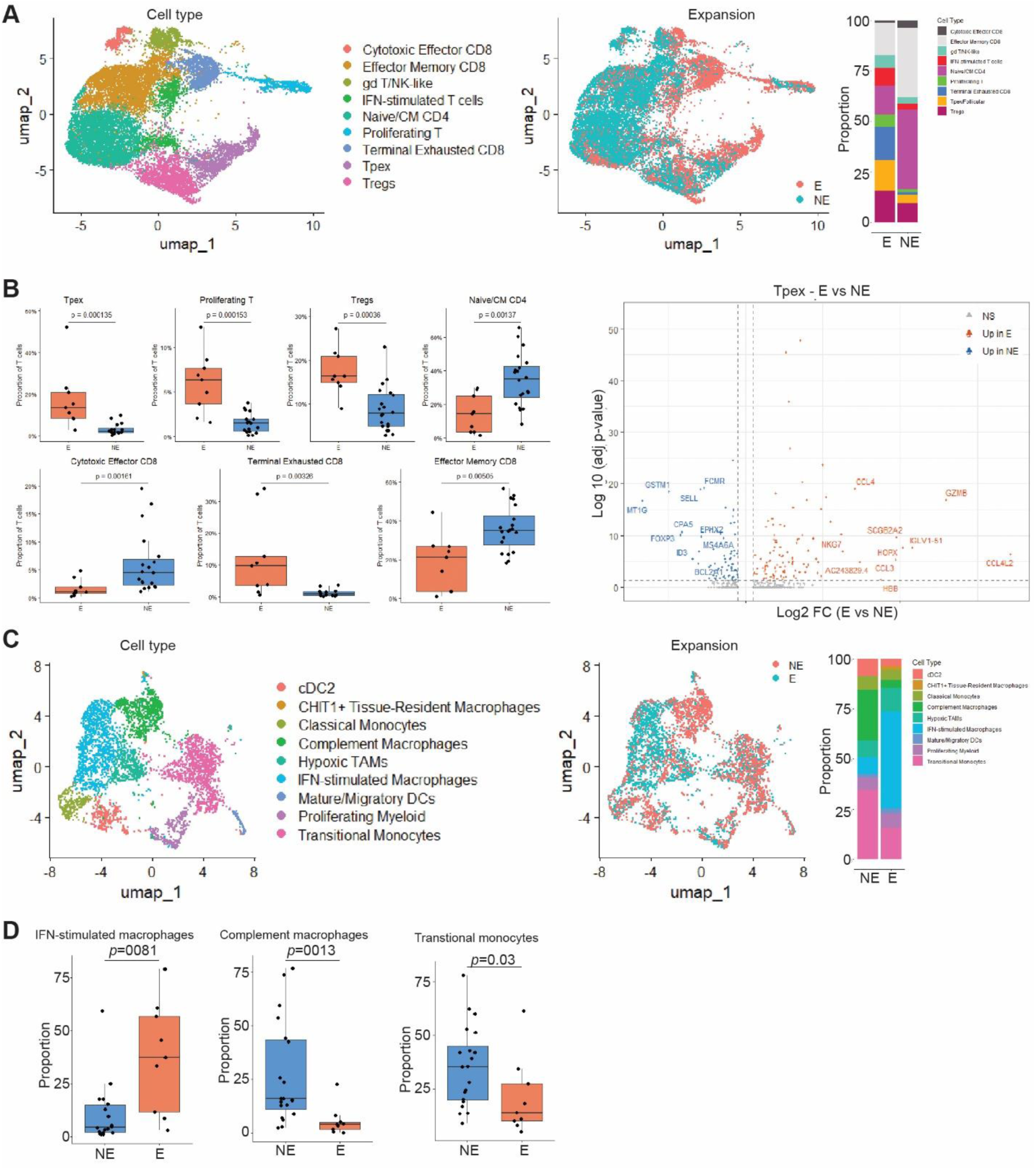
LNP-mIFNα2-treated mouse tumors transcriptionally recapitulate the T cell and myeloid responder phenotypes associated with response to pembrolizumab in human cancer patients. **A.** Human T cell compartment analysis. Left: UMAP of T/NK cells from all patients colored by annotated phenotype, resolving nine distinct subpopulations. Center: same UMAP colored by expansion status (E=orange; NE=blue). Right: Stacked proportion bar chart showing the relative contribution of each T cell phenotype in E vs NE patients. **B.** Quantitative analysis of human T cell subpopulation proportions and differential gene expression. Left: boxplots of comparison of the indicated T cell subpopulations between E (orange) and NE (blue) patients. Right: Volcano plot of differentially expressed genes in the Tpex population comparing E (right, orange) vs. NE (left, blue) patients. **C.** Human myeloid compartment analysis. Left: UMAP of myeloid cells colored by annotated phenotype, resolving nine populations. Center: same UMAP colored by expansion status as indicated. Right: Stacked proportion bar chart comparing myeloid subpopulation composition between E and NE patients. **D.** Boxplot quantification of myeloid subpopulation proportions comparing E (orange) and NE (blue) patients. Boxes indicate median ± interquartile range; whiskers show minima and maxima. Statistical comparisons by Mann-Whitney U test; p-values shown.

In the myeloid compartment, analysis resolved nine macrophage/myeloid phenotypes. E patients were enriched for IFN-stimulated macrophages, while NE patients were dominated by complement macrophages and transitional monocytes (Fig. 7C, D). These findings directly parallel our mouse LNP-mIFNα2 data: the depletion of immunosuppressive TAMs expressing *SPP1*, *ARG1*, and *C1QA*, corresponding to the NE-associated complement macrophage phenotype, and compensatory expansion of Ly6^+^ inflammatory monocytes with ISG signatures mirroring the E-associated IFN-stimulated macrophage phenotype (Fig. 7C, D).

Taken together, these cross-species analyses demonstrate that LNP-mIFNα2 shifts the tumor immune microenvironment into a state transcriptionally and compositionally mirroring the immunotherapy-responsive expander phenotype, expanding cytotoxic and progenitor exhausted T cell subsets while eliminating inhibitory macrophage populations and enriching IFN-stimulated myeloid subsets, thereby establishing a translational link between LNP-mIFNα2-driven reprogramming in mice and effective immunotherapy responses in human cancer.

## DISCUSSION

Most CRC remain refractory to PD-(L)1-based ICI immunotherapy^72^. We show that LNP-mIFNα2 overcomes this resistance through multi-compartment reprogramming of the tumor microenvironment, captured here at single-cell resolution across the myeloid, T, NK, B, and tumor cell compartments. Within the myeloid compartment, LNP-mIFNα2 collapsed immunosuppressive hypoxic SPP1^+^ TAMs and lipid-associated TAMs while expanding inflammatory and patrolling monocytes bearing strong interferon-stimulated gene signatures. We have previously shown that tumor PD-L1 engages myeloid PD-1 to repress IFN-I and impair CXCL9-dependent T cell recruitment^28^ and restoring IFNα2 via lipid nanoparticle delivery of IFNa2 transgene suppress tumor growth^50^. The present mechanistic data indicate that restoring IFN-I directly reverses this circuit, driving repression of SPP1 and apoptotic clearance of SPP1^+^ TAMs rather than their simple reprogramming^73^. The resulting shift away from a complement/SPP1-high macrophage state toward an ISG-high monocyte state directly parallels the myeloid phenotype that distinguishes pembrolizumab responders from non-responders in human breast cancer^71^, reinforcing the myeloid compartment as a primary, rather than bystander, effector of IFNα2-driven tumor growth control.

Two manipulations of the PD-(L)1 axis in this model produced opposite outcomes, and the contrast is instructive rather than contradictory. Anti-PD-1 antibody given before tumor cell inoculation and continued every 3 days thereafter left lung metastatic burden indistinguishable from that of IgG-treated mice, whereas deletion of PD-L1 in the tumor cells themselves nearly abolished lung metastasis. The two manipulations are often read as interchangeable tests of the same axis. They are not. Anti-PD-1 neutralizes only the receptor arm of the interaction, whereas deleting PD-L1 removes the ligand together with every activity of that ligand that PD-1 blockade leaves intact. Tumor PD-L1 also engages CD80, in trans on T cells and in cis on antigen-presenting cells, interactions that constrain T cell priming independently of PD-1 occupancy^31,74^. More pertinent to a therapy built on type I IFN, PD-L1 signals cell-intrinsically within the tumor cell: its conserved intracellular motifs protect tumor cells against IFN-mediated cytotoxicity^75^, and PD-L1 sustains mTOR-dependent glycolytic metabolism and proliferative fitness^76^. CT26.PD-L1 KO cells therefore arrive in the lung stripped of a metabolic and IFN-resistance program that no anti-PD-1 antibody can remove. Genetic deletion is also absolute, uniform across every tumor cell, and operative from the moment of arrival, whereas antibody blockade achieves only partial and fluctuating receptor occupancy, and it does so in a tissue that is only beginning to be infiltrated by the lymphocytes on which the antibody is meant to act.

A further consideration is that ICI releases inhibition and therefore requires a responsive substrate. Our single-cell data show that CD8 *Tpex* cells, the population whose expansion underlies anti-PD-1 efficacy^77^, are scarce in these tumors and held in an *Arg1*/*Txnip*/*Zfp36l2*-high quiescent state. Relieving PD-1 engagement on a numerically insufficient, quiescent progenitor pool accomplishes little. Removing PD-L1 acts further upstream: we have shown previously that tumor PD-L1 engages myeloid PD-1 to repress IFN-I and thereby CXCL9-dependent CTL recruitment to lung metastases^28^, so deleting the ligand lifts that repression constitutively from the moment tumor cells arrive, whereas intermittent receptor blockade acts on a niche in which the IFN-I signal has already been silenced. The failure of anti-PD-1 even when initiated before tumor cells seeded the lung excludes late intervention as the explanation for resistance in this model and places the defect upstream of PD-1 itself. That is the reasoning that led us to supply IFN-I directly, and it predicts, as we observe, that LNP-mIFNα2 restores pharmacologically what PD-L1 deletion restores genetically, and renders these tumors responsive to PD-1 blockade. The mechanism of the PD-L1 knockout phenotype itself was not dissected here; separating loss of tumor cell-intrinsic fitness from restored immune surveillance will require PD-L1 knockout studies in immunodeficient hosts and with lymphocyte depletion, and our deletion is confined to the tumor cell compartment, leaving host PD-L1 intact.

The myeloid remodeling described above was accompanied by relief of CD8 Tpex quiescence: LNP-mIFNα2 reduced expression of the quiescence-associated Zfp36l2, Txnip, and Arg1 program and redirected Tpex cells toward an ISG-high state, with a two-fold expansion of this progenitor-exhausted subset that is mechanistically linked to anti-PD-1 responsiveness^77^. Terminally exhausted CD8 T cells and effector Tregs, which are linked to resistance to PD-1 blockade immunotherapy^78,79^, contracted in parallel, while proliferating and γδ T cells expanded. NK cells showed comparable reprogramming, with both activated and proliferating cytotoxic subsets expanding alongside upregulated cytotoxic effector transcripts, and B cells shifted toward plasma cell and mature states. Together, these compartment-level changes show that IFN-I restoration induces coordinated reprogramming across innate and adaptive immune compartments to convert an exhausted, suppressive infiltrate into a cytolytic one.

A further layer of reprogramming occurred within tumor cells themselves. LNP-mIFNα2 depleted the high-cycling tumor cell state and broadly induced IFN signaling across all tumor subtypes, with consequent upregulation of MHC-I antigen presentation machinery and immune-recruiting chemokines that expanded tumor-immune cell connectivity in CellChat analysis. The IFN-I dependence of this axis is corroborated in pancreatic cancer, where IFNβ mRNA was necessary for cytokine mRNA cocktail-driven MHC-I upregulation and IFNAR neutralization abolished it^44^. Within this same dataset, tumor cells showed a transcriptional pattern consistent with relief of the hypoxia/HIF-1α-driven cuproptosis-resistance program^69^: reduced HIF-1α and glycolytic gene expression, increased copper-importer expression, and reduced copper-sequestering metallothionein, mirroring the reversal of the BACH1-MT1E/X resistance axis in cuproptosis and consistent with the foundational cuproptosis mechanism^67,68,70^. This tumor-intrinsic IFN-cuproptosis crosstalk represents a previously unappreciated layer of LNP-mIFNα2 action that warrants further functional study.

A key mechanistic finding is that antitumor efficacy was substantially dependent on canonical IFNAR1 signaling: anti-IFNAR1 blockade attenuated the effect of LNP-mIFNα2, establishing that efficacy derives from the encoded IFNα2 protein rather than only cGAS-STING sensing of the plasmid backbone^56^. The transient IL6 surge induced by LNP-mIFNα2, a known consequence of cGAS-STING activation^54^, attenuated rather than supported antitumor efficacy. Single-dose IL6 neutralization prior to treatment further reduced metastatic burden, identifying IL6 as a modulatable liability rather than a required component of the antitumor response, and supporting anti-IL6 co-administration as a rational combination strategy.

The LNP-pDNA-associated proinflammatory cytokine and neutrophil profiles remained transient and self-resolved rapidly. The systemic cytokine response was dominated by IFN-I and IFN-driven chemokines rather than broad pro-inflammatory cytokines, and the transient intrapulmonary neutrophil influx resolved fully by 24 hours without tissue injury, including upon repeat boost dosing. This toxicity profile is consistent with findings made in cell-based IFNα2 gene therapy in human cancer patients^43^. A comparable profile, transient cytokine induction and a transient ALT/AST elevation that resolved within 48 h without weight loss, follows systemic nanoparticle delivery of cytokine-encoding mRNAs^44^. Elevated IL6 and GM-CSF in CRC patient serum relative to healthy donors further supports the rationale for IL6 co-blockade in this population.

The premise of this study derives from a clinical result obtained with a different payload. Intismeran autogene, an LNP-delivered mRNA encoding a concatemer of up to 34 patient-specific neoantigens, improved recurrence-free and distant metastasis-free survival when added to pembrolizumab in resected melanoma^6,7^, establishing that an LNP-delivered nucleic acid can change the outcome of checkpoint blockade by supplying an element the tumor microenvironment lacks. We applied the logic of that result rather than its payload. Neoantigen therapy supplies antigen, and therefore presumes both a mutational burden sufficient to nominate immunogenic neoepitopes and an intact antigen-presentation apparatus; neither presumption holds in the pMMR/MSS tumors that constitute 85-90% of CRC. What distinguishes ICI-responsive from ICI-resistant CRC is not antigen supply but a constitutive IFN-high immunophenotype^5^, and IFN-I is the signal that tumor PD-L1 represses within the metastatic niche^28^. We therefore asked whether the missing element could be delivered as the cytokine itself, on the same LNP chassis. The data indicate that it can. A single off-the-shelf formulation controlled metastasis in three PD-1-refractory colon cancer models, including a humanized model of human colon cancer lung metastasis, and did so through IFNAR1, identifying the encoded cytokine rather than innate sensing of the plasmid backbone as the operative agent. Restoring IFN-I did not bypass antigen but restored the means of presenting it: tumor cells induced Tap1/2, Psmb8/9, B2m, and H2-K1/D1 alongside T cell-and NK cell-recruiting chemokines. Consistently, LNP-delivered mRNA encoding no tumor antigen, including SARS-CoV-2 vaccines, sensitizes tumors to checkpoint blockade through early type I IFN induction^19,32^, LNP-mIFNα2 makes that component the payload. Parikh et al. converge from the opposite direction: nanoparticle mRNAs encoding a five-cytokine cocktail, in which IFNβ was required for MHC-I induction, cured autochthonous PDAC without requiring co-delivered antigen^44^. Their cocktail is transient and multiplexed; the nanoplasmid used here sustains a single type I IFN produced by the tumor cells themselves, and in MSS CRC that alone restored antigen presentation and PD-1 responsiveness.

The recent first-in-human clinical trial with Temferon, an engineered autologous HSC product delivering IFNα2 locally within the glioblastoma microenvironment^43^, provides clinical validation of our central premise: tumor-restricted IFN-I reprograms a myeloid-dominant TME without systemic toxicity. It shifted tumor-associated macrophages toward ISG-high inflammatory states, mirroring our findings, but required lentiviral CD34^+^ transduction and myeloablative conditioning. LNP-mIFNα2 achieves convergent multi-compartment reprogramming through a single injection of an off-the-shelf formulation, strengthens the rationale that tumor IFN-I restoration is a generalizable strategy for overcoming myeloid-driven immunotherapy resistance, and supports LNP-IFNα2 for clinical translation in metastatic CRC.

## METHODS

### Cell lines

The murine colon carcinoma cell line CT26 and the human colon adenocarcinoma cell line SW620 were obtained from the American Type Culture Collection (ATCC, Manassas, VA). The MC38-met cell line, a metastatic variant of the MC38 murine colon adenocarcinoma selected for efficient pulmonary colonization, was maintained in culture as previously described^28^. CT26.scramble and CT26.PD-L1 KO cell lines were generated as previously described^28^. All cell lines were cultured in RPMI-1640 supplemented with 10% fetal bovine serum at 37°C in a humidified 5% CO2 incubator. Cell lines were tested bimonthly for mycoplasma and mycoplasma-negative at time of experiments.

### Reagents

The following monoclonal antibodies were used for in vivo treatment: anti-mouse PD-1 (clone RMP1-14, Bio X Cell, Lebanon, NH), Nivolumab (BioXCell), anti-mouse IFNAR1 blocking antibody (clone MAR1-5A3; Bio X Cell), anti-mouse IL6 neutralizing antibody, and corresponding isotype controls (BioXCell).

### LNP-mIFNα2 formulation

A codon-usage-optimized murine IFNα2 (mIFNα2) coding sequence was cloned into the NTC9385R nanoplasmid^80,81^ (Aldevron/Nature Technology Corporation, Fargo, ND). A codon-usage-optimized human IFNα2 (hIFNα2) coding sequence was cloned into NTC9385R. Nanoplasmid DNA was purified in Aldevron under endotoxin-free conditions. Clinical grade DOTAP-cholesterol (1:1) was provided by Genprex Corp (Austin, TX). LNP-mIFNα2 and LNP-hIFNα2 were formulated as previously described ^50^.

### Syngeneic experimental lung metastasis mouse models

CT26 cells (1.5×10^5^ cells/mouse) or MC38-met cells (3.5×10^5^ cells/mouse) were suspended in 100 µl PBS and injected intravenously (i.v.) via the tail vein of female mice on day 0. Mice were randomized into treatment groups on the day of treatment. Treatments consisted of: LNP (control, empty lipid nanoparticles), anti-PD-1 monoclonal antibody (clone RMP1-14, 200 µg/mouse i.p., every 3 days), LNP-mIFNα2 (administered i.v. via tail vein, every 7 days), or LNP-mIFNα2 + anti-PD-1 combination. Two treatment start timepoints were evaluated: day 3 post-tumor injection (early/low burden) and day 5 post-tumor injection (late/high burden). Mice were euthanized at the experimental endpoint, lungs were inflated with ink to visualize tumor nodules as described^82^. All animal experiments were approved by Augusta University and VA Augusta Health Care System Institutional Animal Care and Use Committees (Protocols # 2008-0162 and 22-04-134).

### Human colon cancer cell lung metastasis xenograft humanized mouse model

NOD-scid IL2Rγ-null (NSG) male mice were obtained from The Jackson Laboratory. Humanized mice were generated by intravenous injection of 3×10^6^ human peripheral blood mononuclear cells (PBMCs) isolated from a healthy male donor. Four days after PBMC engraftment, 3×10^6^ SW620 cells were injected i.v. to establish lung metastases. Beginning on Day 5 post-tumor inoculation, mice received LNP control, nivolumab (200 μg/mouse i.p., every 3 days), LNP-hIFNα2 (i.v., every 7 days), or (d) LNP-hIFNα2 + nivolumab combination. Mice were euthanized at the experimental endpoint and lungs were processed for nodule quantification as described above.

### IFNAR1 and IL6 Blockade Experiments

For IFNAR1 dependence studies, CT26 tumor-bearing mice were administered anti-IFNAR1 blocking antibody (200 μg/mouse i.p.) or isotype control IgG prior to LNP-mIFNα2 or LNP-Vector (NCT9385R nanoplasmid) treatment. For IL6 neutralization experiments, anti-IL6 neutralizing monoclonal antibody or isotype control IgG was administered i.p. as a single dose approximately 1h prior to the first LNP-mIFNα2 injection. Pulmonary metastatic nodule counts were assessed at experimental endpoint as the primary efficacy readout.

### Transgene biodistribution and cell-type tropism

To determine the cellular tropism of LNP-mIFNα2 in vivo, CT26 tumor-bearing mice were administered LNP-mIFNα2 i.v. and sacrificed 2 days post-injection. Tumor-bearing lungs were enzymatically digested with collagenase solution and distinct cell populations were isolated by fluorescence-activated cell sorting (FACS) using the following markers: CD45-PD-L1^+^ cells (tumor cells), CD3^+^ cells (T cells), and CD11b^+^ cells (myeloid cells). Sorted populations were lysed and RNA extracted. Transgene expression was detected by PCR using primers specific to the codon-usage-optimized (cuo) mIFNα2 sequence, which is specific for the cuo mIFNα2, not the endogenous mIFNα2^50^. Transgene expression was normalized to the myeloid cell fraction, which was arbitrarily set to 1 as the reference comparator.

### IFNα2 protein pharmacokinetics

To characterize IFNα2 protein production kinetics, CT26 tumor-bearing mice were administered LNP-mIFNα2 i.v. and sacrificed at serial timepoints: 4 hours, 12 hours, and days 1, 2, 3, 6, and 9 post-injection. Blood was collected using a gel column and serum was isolated by centrifugation at 2,000xg for 5 minutes and stored at-80°C until analysis. Tumor-bearing lung tissues were homogenized in total protein lysis buffer with protease inhibitor cocktail, clarified by centrifugation, and protein concentration determined by BCA assay. Serum and lung lysate IFNα2 protein levels were quantified by mouse IFNα2-specific ELISA (Cat# ab211648 Abcam, Cambridge, MA) following the manufacturer’s instructions. This kit specifically detects mouse IFNα2 with no known cross reactivity to other mouse IFNα isoforms. IFNα2 protein level was calculated using R package ^83^ installed on R 3.6.1 (https://www.r-project.org/). The limit of detection was calculated as an average of background samples plus 3 x SD.

### Prime treatment and cytokine analysis

CT26 tumor-bearing mice were treated with LNP, LNP-Vector, or LNP-mIFNα2. Serum was collected 4 hours and 24 hours after treatment. Detection of cytokines in serum or lysates was performed using LEGENDplex Mouse Inflammation Panel (TNFα, IFNγ, IL1α, IL1β, IL6, IL10, IL17A, IL12p70, GM-CSF, IL-23, IFNβ, MCP-1, IL27; BioLegend Cat# 740446), or Mouse Proinflammatory Chemokine Panel V02 (custom 2-plex set for CXCL9 and CXCL10; BioLegend Cat# 741295) according to manufacturer’s instructions. In brief, serum samples were diluted 2-fold with Assay Buffer and standards were mixed with Matrix Solution (Biolegend). For measurements in cell lysates, samples were diluted 4x with Assay Buffer and standards were appropriately mixed with the Lysis Buffer. Standards and samples were plated with capture beads and incubated overnight at 4°C on plate shaker (350 rpm). After washing the plate, detection antibodies were added to each well and the plate was incubated on shaker (600 rpm) for 1h at room temperature. Finally, without washing, SA-PE was added and incubated for 30 min. Samples were acquired on Novocyte Quanteon flow cytometer (Agilent Technologies). Standard curves and protein concentration were calculated using R package ^83^ installed on R 3.6.1. The limit of detection was calculated as an average of background samples plus 3xSD. Total IFNα was measured using Legendplex beads (Biolegend) that detect IFNα1, IFNα6, and IFNα9.

### Boost treatment cytokine analysis

To evaluate the cytokine response upon repeat LNP-mIFNα2 administration, mice that had received a prime LNP-mIFNα2 treatment were re-dosed (boost) with LNP or LNP-mIFNα2 seven days after the prime dose. Serum was collected 4 and 24 hours after the boost treatment and analyzed for the same cytokine/chemokine panel and IFNα2 protein as described for the prime treatment analysis.

### Human colorectal cancer patient serum cytokine analysis

Serum samples were collected from healthy adult donors in Augusta Shepeard Community Blood Center (n=10) and colorectal cancer patients in VA Augusta Health Care System (n=20) under institutional review board (IRB)-approved protocol with written informed consent (IRB Protocol# 1314554-19). The same multiplexed cytokine/chemokine panel used for mouse serum analysis was applied to measure: pro-inflammatory cytokines (IL6, GMCSF, IL-1β, TNFα, IL-8); Th1-associated cytokines (IFNα2, IL12p70, IFNγ, IL2, IFNα, IFNβ, IL28α, IFNL1); Th2-associated cytokines (IL-4, IL-5, IL-9, IL-13); immunoregulatory cytokine IL10; Th17-associated cytokines (IL17A, IL17F, IL22); and chemokines (CXCL10, CXCL5, and an extended chemokine panel). Results were compared between healthy donors and CRC patients by Mann-Whitney U test; p< 0.05 was considered statistically significant.

### Histopathological analysis

Lung tissues from untreated control, LNP-treated, and LNP-mIFNα2-treated CT26 tumor-bearing mice were collected at 4 hours and 24 hours post-treatment. Fixed lungs were processed through graded alcohols, embedded in paraffin, and sectioned at 5 µm thickness. Sections were stained with hematoxylin and eosin (H&E) by standard protocols. H&E-stained sections were evaluated by two board-certified pathologists. The following parameters were assessed: alveolar architecture, evidence of tissue injury (diffuse alveolar damage, necrosis), vascular integrity (vasculitis, endothelial disruption), perivascular inflammation, and intra-alveolar neutrophil content. Quantification of neutrophils was performed by counting neutrophils in 10 consecutive high-power fields (HPF, ×200 magnification) per section; average neutrophil count per 10 HPF was calculated for each animal. Statistical comparisons of neutrophil counts between treatment groups and timepoints were performed by one-way ANOVA with Tukey’s post-hoc test. An analogous histopathological assessment was performed on lung tissues collected 4 and 24 hours after a second (boost) LNP-mIFNα2 dose administered 7 days after the prime treatment.

### Flow cytometry analysis

Tumor-bearing lungs were harvested and enzymatically dissociated using a collagenase solution (1 mg/ml collagenase, Cat#C0130, Collagenase, Sigma-Aldrich, MO; 0.1 mg/ml hyaluronidase, cat#H3506, Sigma-Aldrich; and 30 U/ml DNase I, cat#EN0531, Thermo) at room temperature for 1 h, followed by filtration through a 100 µm cell strainer. Red blood cells were lysed using ACK lysis buffer. Single-cell suspensions were washed in FACS buffer (PBS + 1% FBS). For myeloid infiltrate characterization, cells were stained with the following antibody panels: CD45-FITC, CD11b-APC, Ly6G-PE. For flow cytometric quantification of immune cell subsets (4 hours and 24 hours post-prime and post-boost treatment), cells were acquired on a flow cytometer (BD Fortessa) and data analyzed using FlowJo v10. Gating strategies are shown in Figure 3 and described in detail therein. Cell populations are expressed as percentage of total CD45+ cells, percentage of CD11b+ cells, and as percentage of CD45+CD11b+Ly6G+ fractions. For cell sorting to assess transgene biodistribution, FACS-purified populations (CD45^-^PD-L1^+^, CD3^+^, CD11b^+^) were collected for RNA extraction.

### Single-cell RNA sequencing

CT26 tumor-bearing mice were treated with LNP control or LNP-mIFNα2. At the experimental endpoint, tumor-bearing lungs were harvested and processed to single-cell suspensions as described above. CD45^+^ tumor-infiltrating immune cells were isolated by positive selection using CD45 nanobeads (Biolegend) according to the manufacturer’s protocol, achieving >95% purity as confirmed by post-selection flow cytometry. Cells were resuspended in PBS + 1% BSA. Cell viability was assessed by trypan blue exclusion and was >85% for all samples included in sequencing. Single-cell RNA-seq libraries were generated using the 10x Genomics Chromium platform. Raw BCL files were demultiplexed using Cell Ranger mkfastq v10.0.0. Gene expression was quantified per sample using Cell Ranger count v10.0.0 aligned to the GRCm39-2024-A mouse reference genome. All samples were normalized independently using SCTransform v2 (3,000 variable features, mitochondrial content regressed) and jointly integrated using Reciprocal PCA (RPCA) with SCT normalization across 50 dimensions. PCA was performed on 100 components; the optimal number of PCs(49) was selected using variance-explained criteria. UMAP and tSNE embeddings were computed at 49 PCs. Clustering was performed at resolutions 0.1-1.2; resolution 0.3 yielding 31 clusters was selected as optimal based on Clustree stability analysis. The resulting integrated object served as the sole input for all downstream analysis without re-integration.

### scRNA-sequencing of immune cells

For the immune cell compartment, the integrated object was subsetted to purified CD45^+^ immune cells, preserving existing UMAP coordinates. Unsupervised clustering was performed to identify 31 clusters. Sub-UMAPs, proportion barplots, and count barplots were generated for T cells, NK cells, myeloid cells, and B cells. Cell clusters were annotated to major immune lineages and subpopulations by examination of canonical marker gene expression using DotPlot visualization of known markers. Differentially expressed genes (DEGs) between LNP control and LNP-mIFNα2-treated cells were identified within each annotated cell population using the Wilcoxon rank-sum test implemented in Seurat (FindMarkers), with a minimum log2 fold-change threshold of 0.25 and minimum detection fraction of 0.1 in either group. Adjusted p-values were computed using Bonferroni correction. DEGs are visualized as single-cell heatmaps (log2 fold-change scale). Interferon-stimulated gene (ISG) expression was visualized by DotPlot, with dot size representing percentage of cells expressing each gene and color representing average scaled expression. Cell composition changes were quantified as proportional bar charts and absolute cell counts per annotated population across treatment conditions.

### scRNA sequencing of tumor cell and stromal cells

To collect the non-immune cells, the supernatant fractions of the CD45^+^ cell isolated samples above were loaded on top of lymphocyte separation medium (Cat # 25-072-C1, Mediatech Inc., Manassas, VA) and centrifuged at 2000 RPM for 20 min. The live cells were collected at the interface, washed in PBS, and analyzed for tumor cell/epithelial cell enrichment by flow cytometry. Single-cell library construction and sequencing were performed as described for the CD45^+^ compartment. Raw data were processed using Cell Ranger and analyzed in Seurat using the same pipeline as the immune compartment. The LNP-mIFNα2-treated samples contain only ∼30% CD45^-^ cells, direct comparison of the full LNP-mIFNα2 sample against LNP samples would be confounded by immune cell dilution effects. All CD45^+^ immune contamination is therefore removed via CD45 thresholding before any focused comparison is made. Only after this step are the two samples comparably enriched for CD45^-^ tumor/stromal cells. Supervised clustering resolved seven transcriptionally distinct populations, which were annotated based on canonical mouse colon cancer marker gene expression^64^. Treatment-driven changes in cell population composition were quantified as proportional bar charts and absolute cell counts. DEG analysis between treatment groups within each population was performed as described for the CD45^+^ compartment.

### Cross-species human immunotherapy response analysis

To assess the transcriptomic correspondence between LNP-mIFNα2-induced mouse TME reprogramming and human immunotherapy response signatures, we analyzed a publicly available scRNA-seq dataset from human breast cancer patients receiving neoadjuvant pembrolizumab (EGA study EGAS00001004809, accession#EGAD00001006608)^71^. In this dataset, patients were classified as T cell clonotype Expanders (E) or Non-Expanders (NE) based on T cell receptor clonotype expansion upon pembrolizumab treatment, serving as a surrogate for immunotherapy responsiveness. The human dataset was processed using Seurat with human genome reference GRCh38. Cell type annotation was performed using human-specific canonical markers for T cell subsets (CD8A, CD4, FOXP3, TCF7, HAVCR2, PRF1, GZMB, NKG7, IFNG, CXCL13) and myeloid subsets (CD68, MRC1, C1QA, ISG15, IFITM3, FCN1, CD14). Cell populations were resolved into nine T/NK cell phenotypes and nine myeloid/macrophage phenotypes by UMAP clustering. Proportional composition of each cell population in E versus NE patients was compared using boxplot visualization and Mann-Whitney U test. Differential gene expression in the Tpex/Follicular T cell population between E and NE patients was visualized by volcano plot (log2 fold-change on x-axis,-log10 adjusted *p*-value on y-axis). Gene programs enriched in E patients were compared to gene programs upregulated in LNP-mIFNα2-treated mouse T cell and myeloid populations to identify cross-species transcriptomic concordance.

## Statistical Analysis

All **s**tatistical analysis was carried out using GraphPad Prism 10. Ordinary one-way ANOVA followed by Tukey’s multiple comparisons test and paired Student’s t-test were used to determine statistical significance. For single-cell RNA sequencing data, differential gene expression was assessed by Wilcoxon rank-sum test with Bonferroni correction, and cell subpopulation proportions in the human dataset were compared by Mann-Whitney U test. A *p*<0.05 is considered as significant.

## Acknowledgements

We thank Ms. Jennifer Parks at Augusta Shepeard Community Blood Center for providing blood specimens from healthy donors. We acknowledge the support and contribution of the Integrated Genomics Core Shared Resources at the Georgia Cancer Center, Augusta University (RRID: SCR_026483). We acknowledge the support and contribution of the Immune Monitoring Shared Resource at the Georgia Cancer Center, Augusta University (RRID: SCR_026590), including Rafal Pacholczyk, Valentyna Fesenkova, and Atsuko Matsunaga. We thank Donna Kumiski in the Electron Microscopy & Histology Core Facility at Medical College of Georgia for assistance in electron microscopy and histology.

## Author Contributions

Conceptualization (ZT, PC, PSR, KL), Methodology: (ZT, PC, DBP, HRM, KF, DY), Funding acquisition: (PSR, KL), Project administration (PSR, KL), Bioinformatics (PC, RR, SB), Pathological analysis (JW, LVZ, MZ), Writing: original draft (ZT, PC, PSR, KL), Writing: review & editing: (PC, ZT, JW, LVZ, MZ, PSR, KL)

## Data availability

The scRNA-Seq transcriptomics datasets are deposited in GEO database. Accession #GSE335642. Reviewer token: upsruwsyhjaznun

## REFERENCES

1. Andre, T., Shiu, K.K., Kim, T.W., Jensen, B.V., Jensen, L.H., Punt, C., Smith, D., Garcia-Carbonero, R., Benavides, M., Gibbs, P., et al. (2020). Pembrolizumab in Microsatellite-Instability-High Advanced Colorectal Cancer. N Engl J Med 383, 2207–2218. 10.1056/NEJMoa2017699.

2. Cercek, A., Lumish, M., Sinopoli, J., Weiss, J., Shia, J., Lamendola-Essel, M., El Dika, I.H., Segal, N., Shcherba, M., Sugarman, R., et al. (2022). PD-1 Blockade in Mismatch Repair-Deficient, Locally Advanced Rectal Cancer. N Engl J Med 386, 2363–2376. 10.1056/NEJMoa2201445.

3. Diaz, L.A., Jr., Shiu, K.K., Kim, T.W., Jensen, B.V., Jensen, L.H., Punt, C., Smith, D., Garcia-Carbonero, R., Benavides, M., Gibbs, P., et al. (2022). Pembrolizumab versus chemotherapy for microsatellite instability-high or mismatch repair-deficient metastatic colorectal cancer (KEYNOTE-177): final analysis of a randomised, open-label, phase 3 study. Lancet Oncol 23, 659–670. 10.1016/S1470-2045(22)00197-8.

4. Le, D.T., Kim, T.W., Van Cutsem, E., Geva, R., Jager, D., Hara, H., Burge, M., O’Neil, B., Kavan, P., Yoshino, T., et al. (2020). Phase II Open-Label Study of Pembrolizumab in Treatment-Refractory, Microsatellite Instability-High/Mismatch Repair-Deficient Metastatic Colorectal Cancer: KEYNOTE-164. J Clin Oncol 38, 11–19. 10.1200/JCO.19.02107.

5. Acha-Sagredo, A., Andrei, P., Clayton, K., Taggart, E., Antoniotti, C., Woodman, C.A., Afrache, H., Fourny, C., Armero, M., Moinudeen, H.K., et al. (2025). A constitutive interferon-high immunophenotype defines response to immunotherapy in colorectal cancer. Cancer Cell 43, 292–307 e297. 10.1016/j.ccell.2024.12.008.

6. Fieldhouse, R., and Basu, M. (2026). Moderna cancer vaccine stops melanoma returning: what’s next for personalized treatments? Nature. 10.1038/d41586-026-02612-3.

7. Khattak, A., Carlino, M.S., Meniawy, T., Ansstas, G., Taylor, M.H., Kim, K.B., McKean, M., Long, G.V., Sullivan, R.J., Faries, M., et al. (2026). Intismeran Autogene Plus Pembrolizumab Versus Pembrolizumab Alone in High-Risk Resected Melanoma: 5-Year Update of the Randomized Phase IIb KEYNOTE-942 Study. J Clin Oncol, JCO2600835. 10.1200/JCO-26-00835.

8. Diamond, M.S., Kinder, M., Matsushita, H., Mashayekhi, M., Dunn, G.P., Archambault, J.M., Lee, H., Arthur, C.D., White, J.M., Kalinke, U., et al. (2011). Type I interferon is selectively required by dendritic cells for immune rejection of tumors. J Exp Med 208, 1989– 2003. 10.1084/jem.20101158.

9. Dunn, G.P., Bruce, A.T., Sheehan, K.C., Shankaran, V., Uppaluri, R., Bui, J.D., Diamond, M.S., Koebel, C.M., Arthur, C., White, J.M., and Schreiber, R.D. (2005). A critical function for type I interferons in cancer immunoediting. Nat Immunol 6, 722–729.

10. Shankaran, V., Ikeda, H., Bruce, A.T., White, J.M., Swanson, P.E., Old, L.J., and Schreiber, R.D. (2001). IFNgamma and lymphocytes prevent primary tumour development and shape tumour immunogenicity. Nature 410, 1107–1111. 10.1038/35074122.

11. Zitvogel, L., Galluzzi, L., Kepp, O., Smyth, M.J., and Kroemer, G. (2015). Type I interferons in anticancer immunity. Nat Rev Immunol 15, 405–414. 10.1038/nri3845.

12. Sistigu, A., Yamazaki, T., Vacchelli, E., Chaba, K., Enot, D.P., Adam, J., Vitale, I., Goubar, A., Baracco, E.E., Remedios, C., et al. (2014). Cancer cell-autonomous contribution of type I interferon signaling to the efficacy of chemotherapy. Nat Med 20, 1301–1309. 10.1038/nm.3708nm.3708 [pii].

13. Stone, M.L., Chiappinelli, K.B., Li, H., Murphy, L.M., Travers, M.E., Topper, M.J., Mathios, D., Lim, M., Shih, I.M., Wang, T.L., et al. (2017). Epigenetic therapy activates type I interferon signaling in murine ovarian cancer to reduce immunosuppression and tumor burden. Proc Natl Acad Sci U S A 114, E10981–E10990. 10.1073/pnas.17125141141712514114 [pii].

14. Cauwels, A., Van Lint, S., Garcin, G., Bultinck, J., Paul, F., Gerlo, S., Van der Heyden, J., Bordat, Y., Catteeuw, D., De Cauwer, L., et al. (2018). A safe and highly efficient tumor-targeted type I interferon immunotherapy depends on the tumor microenvironment. Oncoimmunology 7, e1398876. 10.1080/2162402X.2017.13988761398876 [pii].

15. Brown, M.C., Holl, E.K., Boczkowski, D., Dobrikova, E., Mosaheb, M., Chandramohan, V., Bigner, D.D., Gromeier, M., and Nair, S.K. (2017). Cancer immunotherapy with recombinant poliovirus induces IFN-dominant activation of dendritic cells and tumor antigen-specific CTLs. Sci Transl Med 9. eaan4220 [pii]10.1126/scitranslmed.aan42209/408/eaan4220 [pii].

16. Gozgit, J.M., Vasbinder, M.M., Abo, R.P., Kunii, K., Kuplast-Barr, K.G., Gui, B., Lu, A.Z., Molina, J.R., Minissale, E., Swinger, K.K., et al. (2021). PARP7 negatively regulates the type I interferon response in cancer cells and its inhibition triggers antitumor immunity. Cancer Cell 39, 1214–1226 e1210. 10.1016/j.ccell.2021.06.018.

17. Duong, E., Fessenden, T.B., Lutz, E., Dinter, T., Yim, L., Blatt, S., Bhutkar, A., Wittrup, K.D., and Spranger, S. (2021). Type I interferon activates MHC class I-dressed CD11b(+) conventional dendritic cells to promote protective anti-tumor CD8(+) T cell immunity. Immunity. 10.1016/j.immuni.2021.10.020.

18. Propper, D.J., and Balkwill, F.R. (2022). Harnessing cytokines and chemokines for cancer therapy. Nat Rev Clin Oncol. 10.1038/s41571-021-00588-9.

19. Grippin, A.J., Marconi, C., Copling, S., Li, N., Braun, C., Woody, C., Young, E., Gupta, P., Wang, M., Wu, A., et al. (2025). SARS-CoV-2 mRNA vaccines sensitize tumours to immune checkpoint blockade. Nature 647, 488–497. 10.1038/s41586-025-09655-y.

20. Shin, D.S., Zaretsky, J.M., Escuin-Ordinas, H., Garcia-Diaz, A., Hu-Lieskovan, S., Kalbasi, A., Grasso, C.S., Hugo, W., Sandoval, S., Torrejon, D.Y., et al. (2017). Primary Resistance to PD-1 Blockade Mediated by JAK1/2 Mutations. Cancer Discov 7, 188–201. 10.1158/2159-8290.CD-16-1223.

21. Zaretsky, J.M., Garcia-Diaz, A., Shin, D.S., Escuin-Ordinas, H., Hugo, W., Hu-Lieskovan, S., Torrejon, D.Y., Abril-Rodriguez, G., Sandoval, S., Barthly, L., et al. (2016). Mutations Associated with Acquired Resistance to PD-1 Blockade in Melanoma. N Engl J Med 375, 819–829. 10.1056/NEJMoa1604958.

22. Kalbasi, A., Tariveranmoshabad, M., Hakimi, K., Kremer, S., Campbell, K.M., Funes, J.M., Vega-Crespo, A., Parisi, G., Champekar, A., Nguyen, C., et al. (2020). Uncoupling interferon signaling and antigen presentation to overcome immunotherapy resistance due to JAK1 loss in melanoma. Sci Transl Med 12. 10.1126/scitranslmed.abb0152.

23. Torrejon, D.Y., Abril-Rodriguez, G., Champhekar, A.S., Tsoi, J., Campbell, K.M., Kalbasi, A., Parisi, G., Zaretsky, J.M., Garcia-Diaz, A., Puig-Saus, C., et al. (2020). Overcoming Genetically Based Resistance Mechanisms to PD-1 Blockade. Cancer Discov 10, 1140–1157. 10.1158/2159-8290.CD-19-1409.

24. Gao, J., Shi, L.Z., Zhao, H., Chen, J., Xiong, L., He, Q., Chen, T., Roszik, J., Bernatchez, C., Woodman, S.E., et al. (2016). Loss of IFN-gamma Pathway Genes in Tumor Cells as a Mechanism of Resistance to Anti-CTLA-4 Therapy. Cell 167, 397–404 e399. S0092-8674(16)31167-9 [pii]10.1016/j.cell.2016.08.069.

25. Mowat, C., Mosley, S.R., Namdar, A., Schiller, D., and Baker, K. (2021). Anti-tumor immunity in mismatch repair-deficient colorectal cancers requires type I IFN-driven CCL5 and CXCL10. J Exp Med 218. 10.1084/jem.20210108.

26. Huang, K.C., Chiang, S.F., Chang, H.Y., Hong, W.Z., Chen, J.Y., Lee, P.C., Liang, J.A., Ke, T.W., Peng, S.L., Shiau, A.C., et al. (2024). Colorectal cancer-specific IFNbeta delivery overcomes dysfunctional dsRNA-mediated type I interferon signaling to increase the abscopal effect of radiotherapy. J Immunother Cancer 12. 10.1136/jitc-2023-008515.

27. Krieg, A.M. (2025). New insights into the role of IFN-alpha/beta and TLR7/8/9 in cancer immunotherapy and systemic autoimmunity. J Immunother Cancer 13. 10.1136/jitc-2025-012165.

28. Klement, J.D., Redd, P.S., Lu, C., Merting, A.D., Poschel, D.B., Yang, D., Savage, N.M., Zhou, G., Munn, D.H., Fallon, P.G., and Liu, K. (2023). Tumor PD-L1 engages myeloid PD-1 to suppress type I interferon to impair cytotoxic T lymphocyte recruitment. Cancer Cell 41, 620–636 e629. 10.1016/j.ccell.2023.02.005.

29. Hui, E., Cheung, J., Zhu, J., Su, X., Taylor, M.J., Wallweber, H.A., Sasmal, D.K., Huang, J., Kim, J.M., Mellman, I., and Vale, R.D. (2017). T cell costimulatory receptor CD28 is a primary target for PD-1-mediated inhibition. Science 355, 1428–1433. 10.1126/science.aaf1292.

30. Kamphorst, A.O., Wieland, A., Nasti, T., Yang, S., Zhang, R., Barber, D.L., Konieczny, B.T., Daugherty, C.Z., Koenig, L., Yu, K., et al. (2017). Rescue of exhausted CD8 T cells by PD-1-targeted therapies is CD28-dependent. Science 355, 1423–1427. 10.1126/science.aaf0683science.aaf0683 [pii].

31. Sugiura, D., Maruhashi, T., Okazaki, I.M., Shimizu, K., Maeda, T.K., Takemoto, T., and Okazaki, T. (2019). Restriction of PD-1 function by cis-PD-L1/CD80 interactions is required for optimal T cell responses. Science 364, 558–566. 10.1126/science.aav7062.

32. Qdaisat, S., Wummer, B., Stover, B.D., Zhang, D., McGuiness, J., Weidert, F., Chardon-Robles, J., Grippin, A., DeVries, A., Zhao, C., et al. (2025). Sensitization of tumours to immunotherapy by boosting early type-I interferon responses enables epitope spreading. Nat Biomed Eng 9, 1437–1452. 10.1038/s41551-025-01380-1.

33. Kirkwood, J. (2002). Cancer immunotherapy: the interferon-alpha experience. Semin Oncol 29, 18–26. S0093-7754(02)50195-0 [pii].

34. Garbe, C., and Eigentler, T.K. (2007). Diagnosis and treatment of cutaneous melanoma: state of the art 2006. Melanoma Res 17, 117–127. 10.1097/CMR.0b013e328042bb3600008390-200704000-00007 [pii].

35. Hervas-Stubbs, S., Perez-Gracia, J.L., Rouzaut, A., Sanmamed, M.F., Le Bon, A., and Melero, I. (2011). Direct effects of type I interferons on cells of the immune system. Clin Cancer Res 17, 2619–2627. 10.1158/1078-0432.CCR-10-11141078-0432.CCR-10-1114 [pii].

36. Trinchieri, G. (2010). Type I interferon: friend or foe? J Exp Med 207, 2053–2063. 10.1084/jem.20101664.

37. Hauschild, A., Dummer, R., Ugurel, S., Kaehler, K.C., Egberts, F., Fink, W., Both-Skalsky, J., Laetsch, B., and Schadendorf, D. (2008). Combined treatment with pegylated interferon-alpha-2a and dacarbazine in patients with advanced metastatic melanoma: a phase 2 study. Cancer 113, 1404–1411. 10.1002/cncr.23722.

38. Eliason, J.F. (2001). Pegylated cytokines: potential application in immunotherapy of cancer. BioDrugs 15, 705–711. 10.2165/00063030-200115110-00001.

39. Lee, S., and Margolin, K. (2011). Cytokines in cancer immunotherapy. Cancers (Basel) 3, 3856–3893. 10.3390/cancers3043856.

40. Dummer, R., and Mangana, J. (2009). Long-term pegylated interferon-alpha and its potential in the treatment of melanoma. Biologics 3, 169–182. 10.2147/btt.2009.3051.

41. Cao, X., Liang, Y., Hu, Z., Li, H., Yang, J., Hsu, E.J., Zhu, J., Zhou, J., and Fu, Y.X. (2021). Next generation of tumor-activating type I IFN enhances anti-tumor immune responses to overcome therapy resistance. Nat Commun 12, 5866. 10.1038/s41467-021-26112-2.

42. Yang, X., Zhang, X., Fu, M.L., Weichselbaum, R.R., Gajewski, T.F., Guo, Y., and Fu, Y.X. (2014). Targeting the tumor microenvironment with interferon-beta bridges innate and adaptive immune responses. Cancer Cell 25, 37–48. 10.1016/j.ccr.2013.12.004.

43. Gentner, B., Eoli, M., Farina, F., Barcella, M., Capotondo, A., Mazzoleni, S., Brambilla, V., Francavilla, A., Garramone, M., Carrabba, M.G., et al. (2026). Tumor-targeted interferon-alpha gene therapy for glioblastoma: a phase 1 trial. Nat Med 32, 2216–2226. 10.1038/s41591-026-04419-1.

44. Parikh, C.N., DeMarco, K.D., Bhalerao, N., Giwa, H.K., Kane, G.I., Dinnell, R.W., Ma, B., Mori, H., Brassil, M.L., Murphy, K.C., et al. (2026). Multiplexed cytokine and antigen mRNA administration generates durable anti-tumor immunity against pancreatic cancer. Nat Commun 17. 10.1038/s41467-026-74574-z.

45. Templeton, N.S., Lasic, D.D., Frederik, P.M., Strey, H.H., Roberts, D.D., and Pavlakis, G.N. (1997). Improved DNA: liposome complexes for increased systemic delivery and gene expression. Nat Biotechnol 15, 647–652. 10.1038/nbt0797-647.

46. Lu, C., Stewart, D.J., Lee, J.J., Ji, L., Ramesh, R., Jayachandran, G., Nunez, M.I., Wistuba, II, Erasmus, J.J., Hicks, M.E., et al. (2012). Phase I clinical trial of systemically administered TUSC2(FUS1)-nanoparticles mediating functional gene transfer in humans. PLoS One 7, e34833. 10.1371/journal.pone.0034833.

47. Meraz, I.M., Majidi, M., Cao, X., Lin, H., Li, L., Wang, J., Baladandayuthapani, V., Rice, D., Sepesi, B., Ji, L., and Roth, J.A. (2018). TUSC2 Immunogene Therapy Synergizes with Anti-PD-1 through Enhanced Proliferation and Infiltration of Natural Killer Cells in Syngeneic Kras-Mutant Mouse Lung Cancer Models. Cancer Immunol Res 6, 163–177. 10.1158/2326-6066.CIR-17-0273.

48. Al Subeh, Z.Y., Poschel, D.B., Redd, P.S., Klement, J.D., Merting, A.D., Yang, D., Mehta, M., Shi, H., Colson, Y.L., Oberlies, N.H., et al. (2022). Lipid Nanoparticle Delivery of Fas Plasmid Restores Fas Expression to Suppress Melanoma Growth In Vivo. ACS Nano 16, 12695–12710. 10.1021/acsnano.2c04420.

49. Merting, A.D., Poschel, D.B., Lu, C., Klement, J.D., Yang, D., Li, H., Shi, H., Chapdelaine, E., Montgomery, M., Redman, M.T., et al. (2022). Restoring FAS Expression via Lipid-Encapsulated FAS DNA Nanoparticle Delivery Is Sufficient to Suppress Colon Tumor Growth In Vivo. Cancers (Basel) 14. 10.3390/cancers14020361.

50. Fick, K., Kerns, N., Zhao, Y., Tiamiyu, Z., Poschel, D., Czabala, P., Yang, D., Tang, Y., Xie, J., Fesenkova, V., et al. (2025). Lipid nanoparticle-delivered IFNalpha2 activates Cxcl9 to increase T cell tumor recruitment to suppress lung metastasis. J Immunother Cancer 13. 10.1136/jitc-2024-011415.

51. Seyed, N., Zahedifard, F., Habibzadeh, S., Yousefi, R., Lajevardi, M.S., Gholami, E., and Rafati, S. (2022). Antibiotic-Free Nanoplasmids as Promising Alternatives for Conventional DNA Vectors. Vaccines (Basel) 10. 10.3390/vaccines10101710.

52. Liu, S.Y., Huang, W.C., Yeh, H.I., Ko, C.C., Shieh, H.R., Hung, C.L., Chen, T.Y., and Chen, Y.J. (2019). Sequential Blockade of PD-1 and PD-L1 Causes Fulminant Cardiotoxicity-From Case Report to Mouse Model Validation. Cancers (Basel) 11. 10.3390/cancers11040580.

53. Song, W., Shen, L., Wang, Y., Liu, Q., Goodwin, T.J., Li, J., Dorosheva, O., Liu, T., Liu, R., and Huang, L. (2018). Synergistic and low adverse effect cancer immunotherapy by immunogenic chemotherapy and locally expressed PD-L1 trap. Nat Commun 9, 2237. 10.1038/s41467-018-04605-x.

54. Patel, M.N., Tiwari, S., Wang, Y., O’Neill, S., Wu, J., Omo-Lamai, S., Espy, C., Chase, L.S., Majumder, A., Hoffman, E., et al. (2026). Safer non-viral DNA delivery using lipid nanoparticles loaded with endogenous anti-inflammatory lipids. Nat Biotechnol 44, 79–89. 10.1038/s41587-025-02556-5.

55. Schmitt, M., and Greten, F.R. (2021). The inflammatory pathogenesis of colorectal cancer. Nat Rev Immunol 21, 653–667. 10.1038/s41577-021-00534-x.

56. Negishi, H., Wada, Y., Shirasaki, Y., Hayashi, T., Kubota, Y., Iwasaki, T., Kurosawa, M., Ban, T., Muto, D., Suenaga, Y., et al. (2026). cGAS-IFN-I responses by extracting nuclear DNA from dying cells via nucleocytosis. Nat Commun 17, 1658. 10.1038/s41467-026-68839-w.

57. Hailemichael, Y., Johnson, D.H., Abdel-Wahab, N., Foo, W.C., Bentebibel, S.E., Daher, M., Haymaker, C., Wani, K., Saberian, C., Ogata, D., et al. (2022). Interleukin-6 blockade abrogates immunotherapy toxicity and promotes tumor immunity. Cancer Cell 40, 509–523 e506. 10.1016/j.ccell.2022.04.004.

58. Liu, S., Zhang, Z., Wang, Z., Liu, C., Liang, G., Xu, T., Li, Z., Duan, X., Xu, G., Feng, X., et al. (2026). SPP1 Drives Colorectal Cancer Liver Metastasis and Immunotherapy Resistance by Stimulating CXCL12 Production in Cancer-Associated Fibroblasts. Cancer Res 86, 58– 79. 10.1158/0008-5472.CAN-24-4916.

59. Qi, J., Sun, H., Zhang, Y., Wang, Z., Xun, Z., Li, Z., Ding, X., Bao, R., Hong, L., Jia, W., et al. (2022). Single-cell and spatial analysis reveal interaction of FAP(+) fibroblasts and SPP1(+) macrophages in colorectal cancer. Nat Commun 13, 1742. 10.1038/s41467-022-29366-6.

60. Trehan, R., Huang, P., Zhu, X.B., Wang, X., Soliman, M., Strepay, D., Nur, A., Kedei, N., Arhin, M., Ghabra, S., et al. (2025). SPP1 + macrophages cause exhaustion of tumor-specific T cells in liver metastases. Nat Commun 16, 4242. 10.1038/s41467-025-59529-0.

61. Bill, R., Wirapati, P., Messemaker, M., Roh, W., Zitti, B., Duval, F., Kiss, M., Park, J.C., Saal, T.M., Hoelzl, J., et al. (2023). CXCL9:SPP1 macrophage polarity identifies a network of cellular programs that control human cancers. Science 381, 515–524. 10.1126/science.ade2292.

62. Kwart, D., He, J., Srivatsan, S., Lett, C., Golubov, J., Oswald, E.M., Poon, P., Ye, X., Waite, J., Zaretsky, A.G., et al. (2022). Cancer cell-derived type I interferons instruct tumor monocyte polarization. Cell Rep 41, 111769. 10.1016/j.celrep.2022.111769.

63. Elewaut, A., Estivill, G., Bayerl, F., Castillon, L., Novatchkova, M., Pottendorfer, E., Hoffmann-Haas, L., Schonlein, M., Nguyen, T.V., Lauss, M., et al. (2025). Cancer cells impair monocyte-mediated T cell stimulation to evade immunity. Nature 637, 716–725. 10.1038/s41586-024-08257-4.

64. Gao, J., Wu, Z., Zhao, M., Zhang, R., Li, M., Sun, D., Cheng, H., Qi, X., Shen, Y., Xu, Q., et al. (2022). Allosteric inhibition reveals SHP2-mediated tumor immunosuppression in colon cancer by single-cell transcriptomics. Acta Pharm Sin B 12, 149–166. 10.1016/j.apsb.2021.08.006.

65. Ahmed, M., Sottnik, J.L., Dancik, G.M., Sahu, D., Hansel, D.E., Theodorescu, D., and Schwartz, M.A. (2016). An Osteopontin/CD44 Axis in RhoGDI2-Mediated Metastasis Suppression. Cancer Cell 30, 432–443. 10.1016/j.ccell.2016.08.002.

66. Czabala, P., Zhao, Y., Klement, J.D., Redd, P.S., Poschel, D., Carver, K., Fick, K., Tiamiyu, Z., Zoccheddu, M., Schoenlein, P., et al. (2026). Distinct and cooperative roles of host and tumor Osteopontin in colorectal cancer liver metastasis. bioRxiv, 2026.2002.2019.706899. 10.64898/2026.02.19.706899.

67. Lei, G., Lu, Z., Xu, Z., Braun, C., Huo, D., Gao, J., Tan, L., Hong, T., Wu, S., Sun, M., et al. (2026). Cuproptosis-immunity crosstalk informs strategy to overcome immunotherapy resistance. Cell. 10.1016/j.cell.2026.05.036.

68. Lei, G., Sun, M., Cheng, J., Ye, R., Lu, Z., Horbath, A., Huo, D., Wu, S., Alapati, A., Aggarwal, S., et al. (2025). Radiotherapy promotes cuproptosis and synergizes with cuproptosis inducers to overcome tumor radioresistance. Cancer Cell 43, 1076–1092 e1075. 10.1016/j.ccell.2025.03.031.

69. Yang, Z., Su, W., Wei, X., Pan, Y., Xing, M., Niu, L., Feng, B., Kong, W., Ren, X., Huang, F., et al. (2025). Hypoxia inducible factor-1alpha drives cancer resistance to cuproptosis. Cancer Cell 43, 937–954 e939. 10.1016/j.ccell.2025.02.015.

70. Dang, D., Deogharkar, A., McKolay, J., Smith, K.S., Panwalkar, P., Hoffman, S., Tian, W., Ji, S., Azambuja, A.P., Natarajan, S.K., et al. (2025). Isocitrate dehydrogenase 1 primes group-3 medulloblastomas for cuproptosis. Cancer Cell 43, 1159–1174 e1158. 10.1016/j.ccell.2025.04.013.

71. Bassez, A., Vos, H., Van Dyck, L., Floris, G., Arijs, I., Desmedt, C., Boeckx, B., Vanden Bempt, M., Nevelsteen, I., Lambein, K., et al. (2021). A single-cell map of intratumoral changes during anti-PD1 treatment of patients with breast cancer. Nat Med 27, 820–832. 10.1038/s41591-021-01323-8.

72. Brahmer, J.R., Tykodi, S.S., Chow, L.Q., Hwu, W.J., Topalian, S.L., Hwu, P., Drake, C.G., Camacho, L.H., Kauh, J., Odunsi, K., et al. (2012). Safety and activity of anti-PD-L1 antibody in patients with advanced cancer. N Engl J Med 366, 2455–2465. 10.1056/NEJMoa1200694.

73. Wei, J., Marisetty, A., Schrand, B., Gabrusiewicz, K., Hashimoto, Y., Ott, M., Grami, Z., Kong, L.Y., Ling, X., Caruso, H., et al. (2019). Osteopontin mediates glioblastoma-associated macrophage infiltration and is a potential therapeutic target. J Clin Invest 129, 137–149. 10.1172/JCI121266.

74. Butte, M.J., Keir, M.E., Phamduy, T.B., Sharpe, A.H., and Freeman, G.J. (2007). Programmed death-1 ligand 1 interacts specifically with the B7-1 costimulatory molecule to inhibit T cell responses. Immunity 27, 111–122. 10.1016/j.immuni.2007.05.016.

75. Gato-Canas, M., Zuazo, M., Arasanz, H., Ibanez-Vea, M., Lorenzo, L., Fernandez-Hinojal, G., Vera, R., Smerdou, C., Martisova, E., Arozarena, I., et al. (2017). PDL1 Signals through Conserved Sequence Motifs to Overcome Interferon-Mediated Cytotoxicity. Cell Rep 20, 1818–1829. 10.1016/j.celrep.2017.07.075.

76. Chang, C.H., Qiu, J., O’Sullivan, D., Buck, M.D., Noguchi, T., Curtis, J.D., Chen, Q., Gindin, M., Gubin, M.M., van der Windt, G.J., et al. (2015). Metabolic Competition in the Tumor Microenvironment Is a Driver of Cancer Progression. Cell 162, 1229–1241. 10.1016/j.cell.2015.08.016.

77. Nose, Y., Yasumizu, Y., Saito, T., Nakamura, Y., Jinushi, K., Fujikawa, K., Momose, K., Yamashita, K., Tanaka, K., Yamamoto, K., et al. (2026). PD-1 antibody-bound progenitor-exhausted CD8(+) T cells in lymph nodes boost PD-1-blockade anti-tumor immunity in gastrointestinal cancer. Nat Commun 17. 10.1038/s41467-026-70751-2.

78. Chakravarti, M., Dhar, S., Bera, S., Sinha, A., Roy, K., Sarkar, A., Dasgupta, S., Bhuniya, A., Saha, A., Das, J., et al. (2023). Terminally Exhausted CD8+ T Cells Resistant to PD-1 Blockade Promote Generation and Maintenance of Aggressive Cancer Stem Cells. Cancer Res 83, 1815–1833. 10.1158/0008-5472.CAN-22-3864.

79. Watson, M.J., Vignali, P.D.A., Mullett, S.J., Overacre-Delgoffe, A.E., Peralta, R.M., Grebinoski, S., Menk, A.V., Rittenhouse, N.L., DePeaux, K., Whetstone, R.D., et al. (2021). Metabolic support of tumour-infiltrating regulatory T cells by lactic acid. Nature 591, 645– 651. 10.1038/s41586-020-03045-2.

80. Souto, E.B., Blanco-Llamero, C., Krambeck, K., Kiran, N.S., Yashaswini, C., Postwala, H., Severino, P., Priefer, R., Prajapati, B.G., and Maheshwari, R. (2024). Regulatory insights into nanomedicine and gene vaccine innovation: Safety assessment, challenges, and regulatory perspectives. Acta Biomater 180, 1–17. 10.1016/j.actbio.2024.04.010.

81. Williams, J.A. (2013). Vector Design for Improved DNA Vaccine Efficacy, Safety and Production. Vaccines (Basel) 1, 225–249. 10.3390/vaccines1030225.

82. Paschall, A.V., and Liu, K. (2016). An Orthotopic Mouse Model of Spontaneous Breast Cancer Metastasis. J Vis Exp. 10.3791/54040.

83. Sanz, H., Aponte, J.J., Harezlak, J., Dong, Y., Ayestaran, A., Nhabomba, A., Mpina, M., Maurin, O.R., Diez-Padrisa, N., Aguilar, R., et al. (2017). drLumi: An open-source package to manage data, calibrate, and conduct quality control of multiplex bead-based immunoassays data analysis. PLoS One 12, e0187901. 10.1371/journal.pone.0187901.

