## Supplemental data for "Multi-compartment immune and tumor cell reprogramming by LNP-IFNα2 overcomes colon cancer immunotherapy resistance"

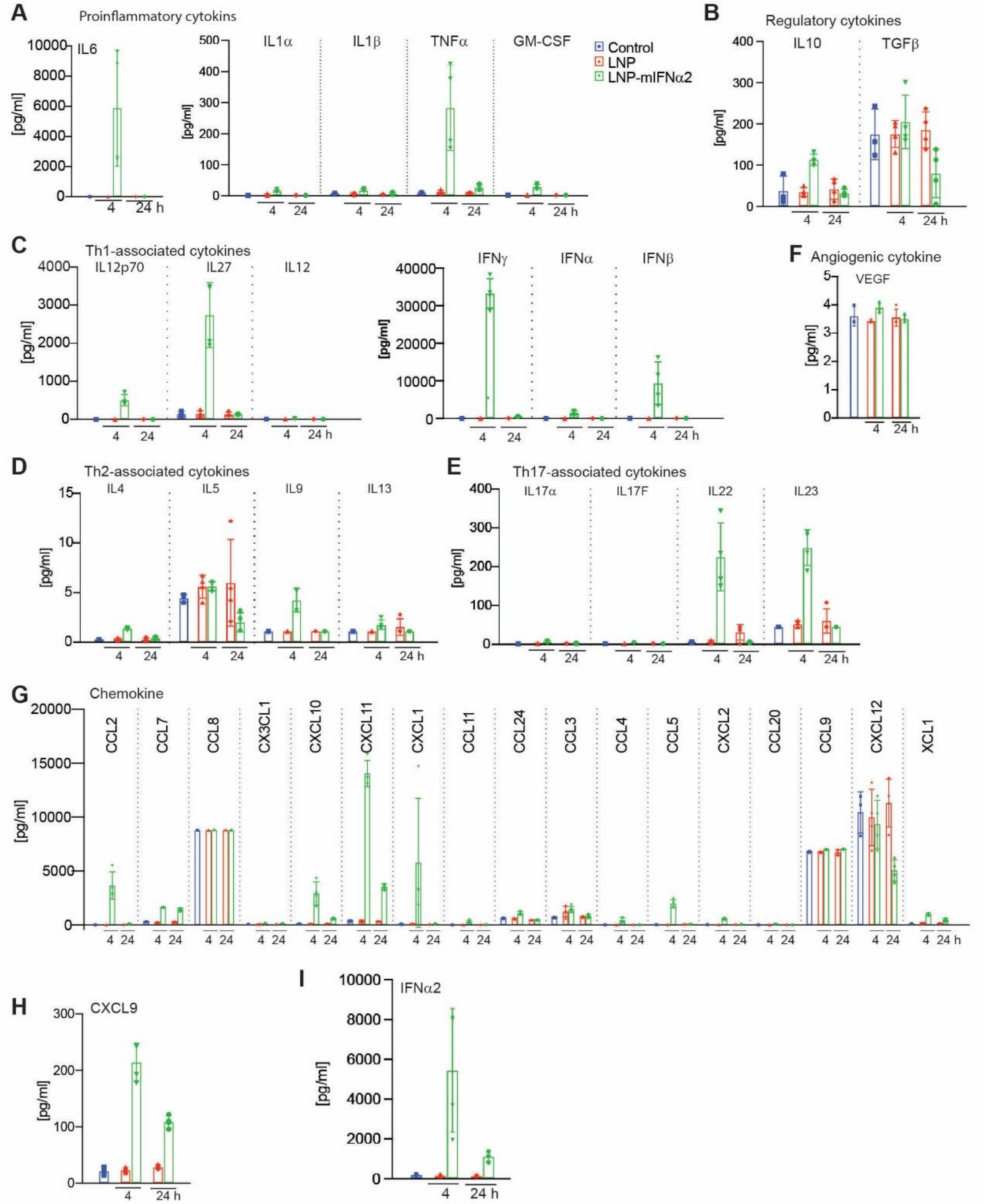


+

**Figure S1. LNP-mIFNα2 boost therapy induces a transient type I interferon and chemokine response. A**. Pro-inflammatory cytokines in mouse serum. The LNP-mIFNα2-treated mice were treated again 7 days later with LNP or LNP-mIFNα2. Serum was then collected 4 and 24h after the boost treatment. Data represents mean±SD. **B-F**. Serum cytokines from mice at 4 and 24h after the LNP or LNP-mIFNα2 boost treatment are measured by multiplex immunoassay as in A and grouped by functional class as indicated. **G-H**. Chemokines from mice at 4 and 24h after LNP or LNP-mIFNα2 boost treatment are measured by multiplex immunoassay as in A. **I**. Serum IFNα2 protein levels from mice at 4 and 24h after LNP or LNP-mIFNα2 boost treatment were measured by mouse IFNα2-specific ELISA.

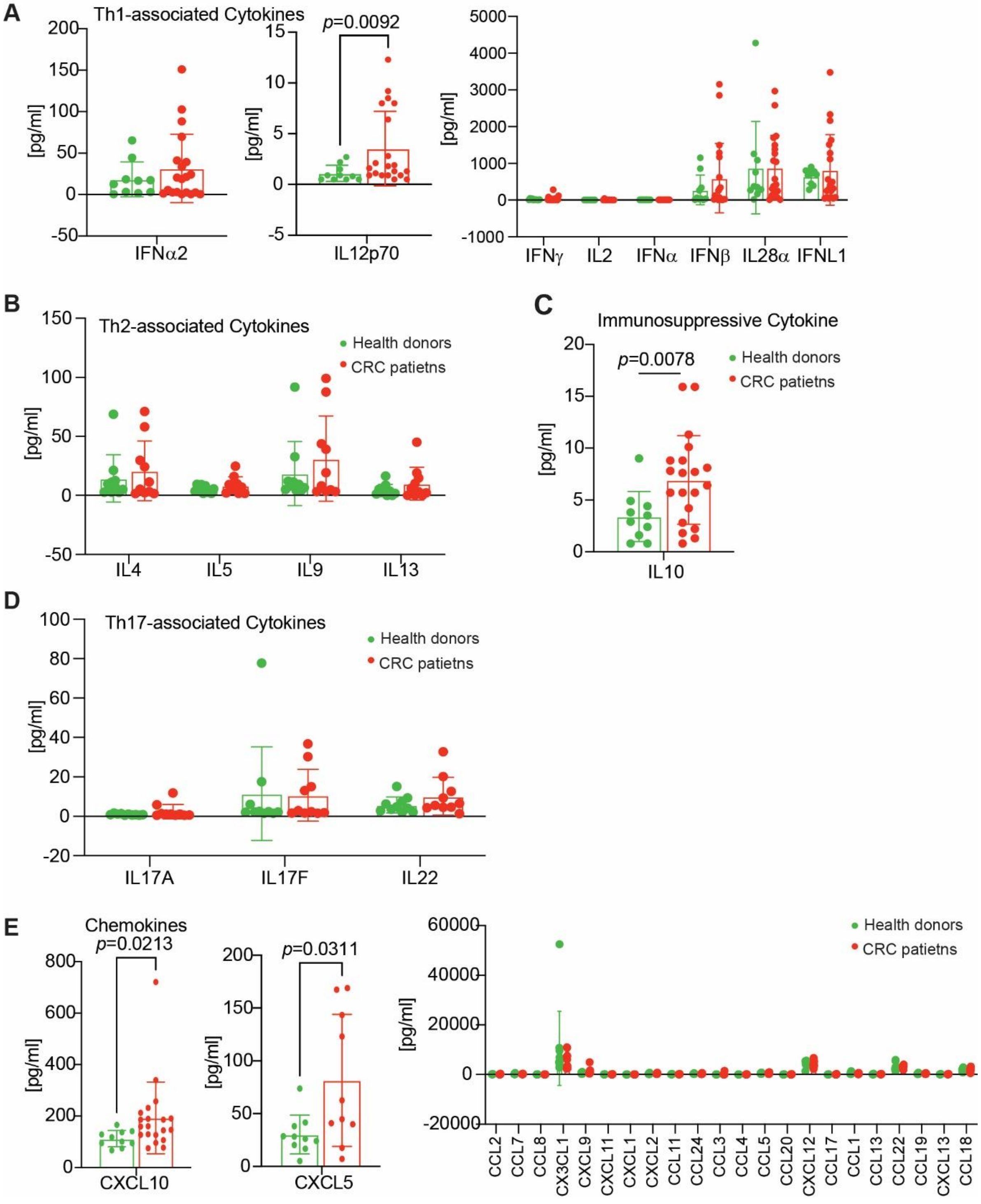


**Figure S2. Serum cytokine and chemokine profiles in colorectal cancer patients versus healthy donors. A.** Th1-associated cytokines in serum from healthy donors (green) and CRC patients (red). **B.** Th2-associated cytokines. **C.** Immunoregulatory cytokine IL-10. **D.** Th17-

associated cytokines. **E.** Chemokines. Each point represents one donor or patient. Data represents mean±SD. Only significant differences are noted.

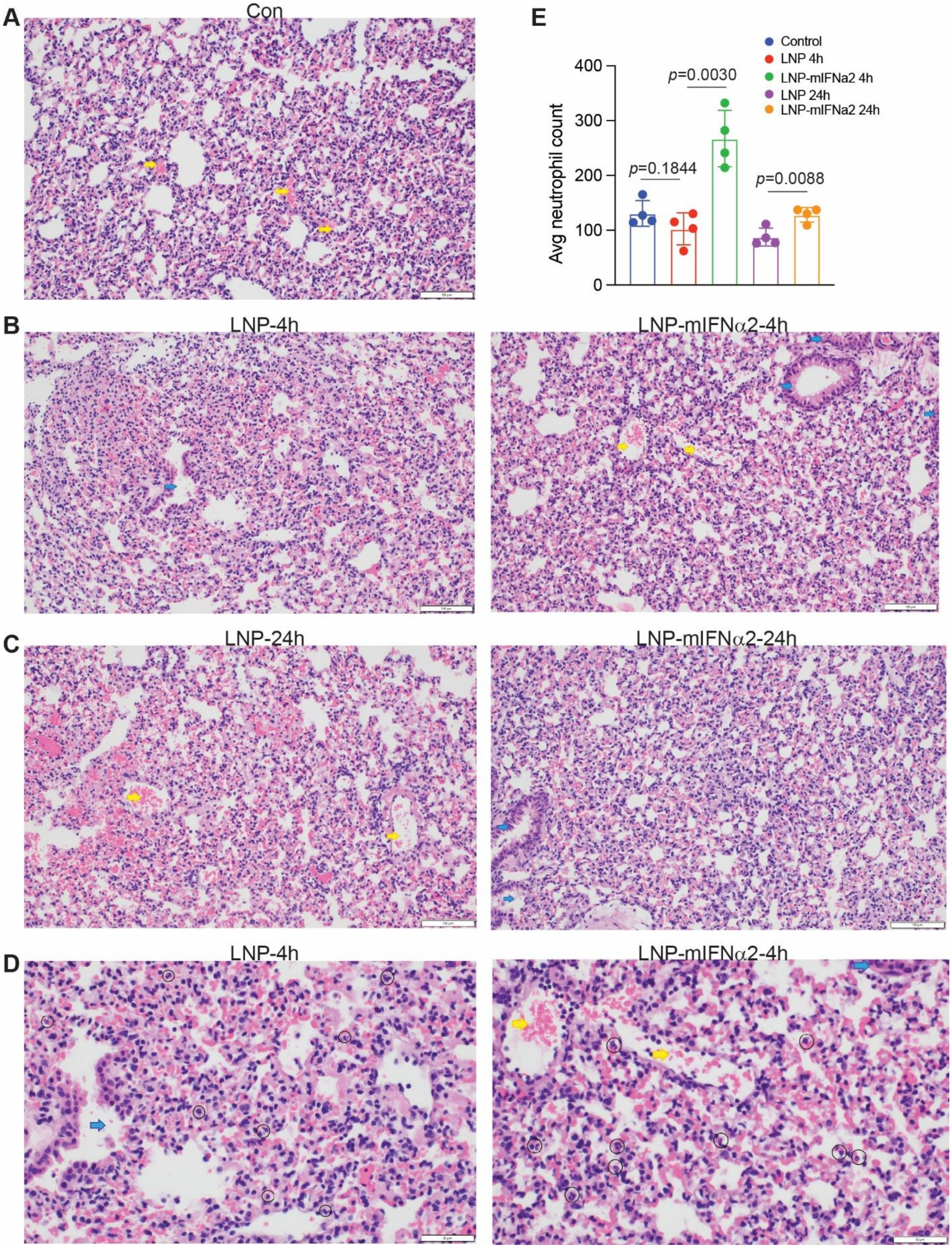

**Figure S3. LNP-mIFNα2 induces a transient and self-resolving alveolar neutrophil influx. A.** Representative H&E-stained lung section from an untreated control CT26 metastasis-bearing mouse. Yellow arrow: Pulmonary arteries; Blue arrow: bronchioles. **B-C.** Representative H&E-stained lung sections at 4 (B) and 24 (C) hours after LNP (left) and LNP-mIFNα2 (right) treatment. **D.** Higher-magnification H&E images of representative lung sections from LNP-treated (left) and LNP-mIFNα2-treated (right) mice at 4 h post-treatment. Black circles indicate neutrophils. Blue arrow: bronchiole. Yellow arrow: pulmonary artery. The vessel walls remained intact, with no evidence of vasculitis or endothelial disruption. Circle: neutrophils located within the alveolar space. **E.** Quantification of average intra-alveolar neutrophil counts per 10 HPF across all experimental groups as indicated. Each data point represents one mouse (n=4 per group). Data represent mean±SD.

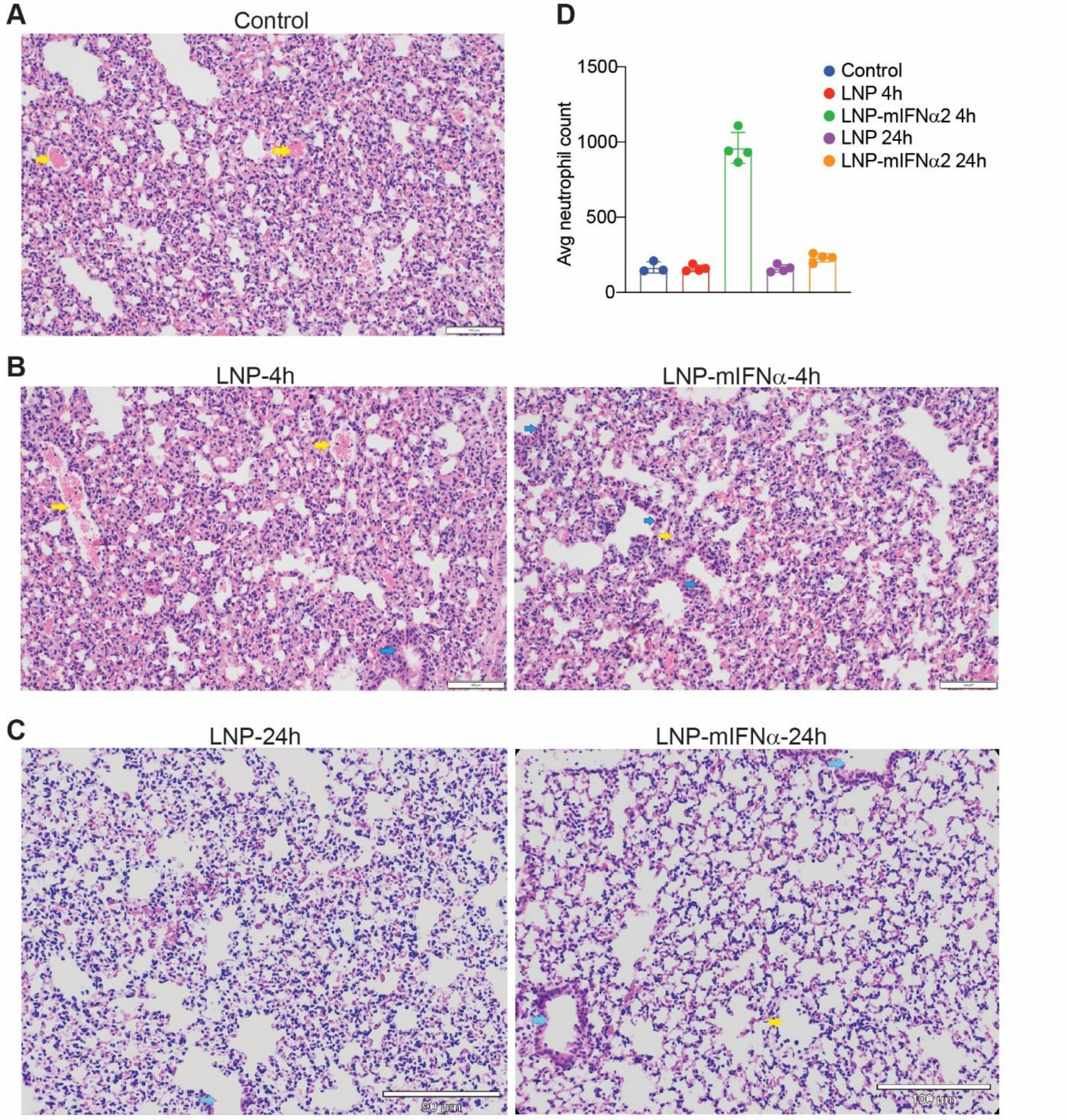


**Figure S4. LNP-mIFNa2 boost therapy also induces a transient and self-resolving alveolar neutrophil influx. A.** Representative H&E-stained lung section from the LNP-mIFNα2-treated mice at day 7 without further treatment and used as control for the boost treatment. Yellow arrow: Pulmonary arteries; Blue arrow: bronchioles. **B-C.** Representative H&E-stained lung sections at 4 (B) and 24 (C) hours after LNP (left) and LNP-mIFNα2 (right) boost treatment 7 days after the prime treatment with LNP-mIFNα2. **D**. Quantification of average intra-alveolar neutrophil counts per 10 HPF across all experimental groups as indicated. Each data point represents one mouse. Data represent mean±SD.

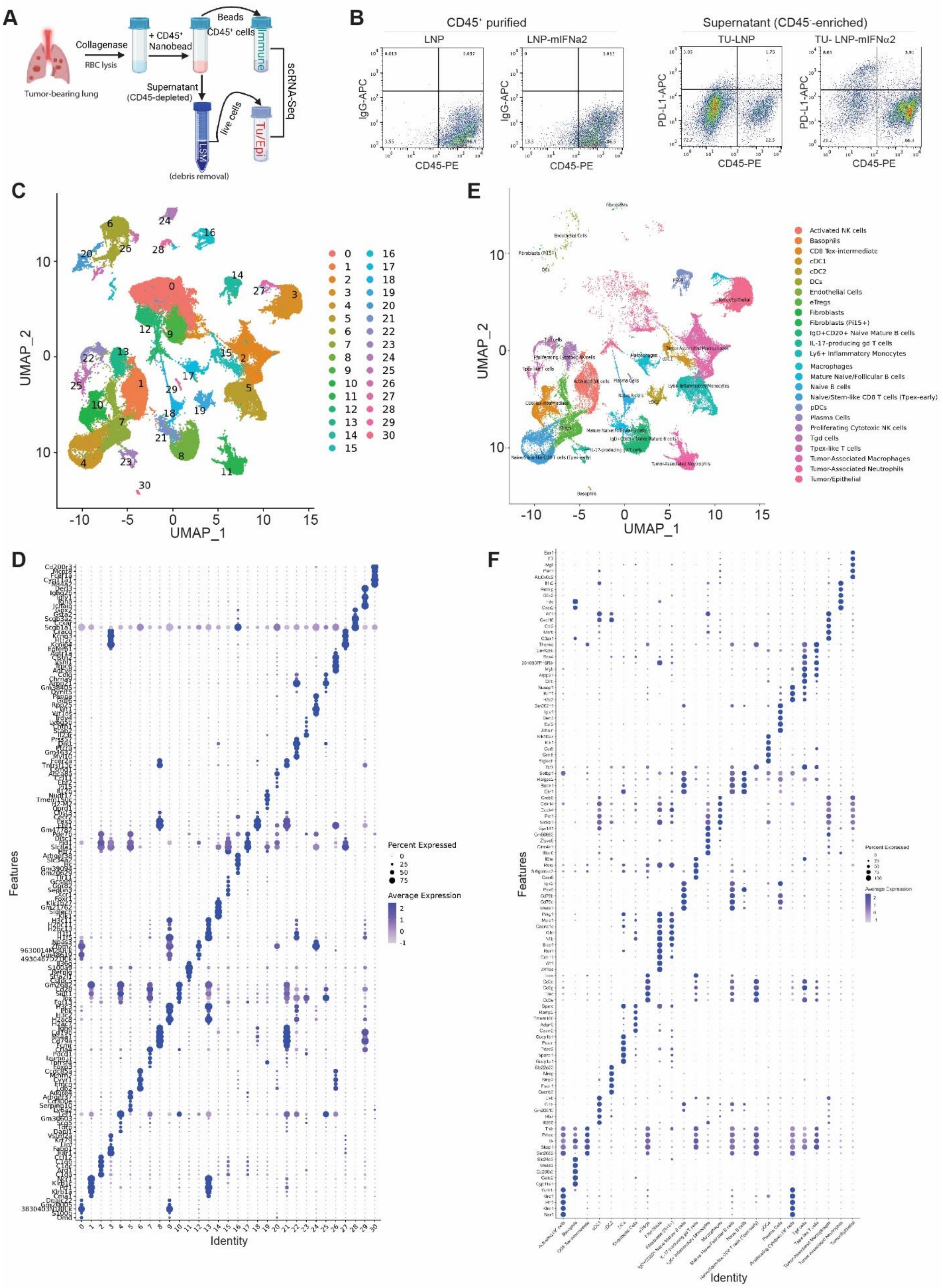
A
Tumor-bearing lung
Collagenase
RBC lysis
+ CD45+ Nanobead
Beads
CD45+ cells
Immune
scRNA-Seq
Supernatant (CD45-depleted)
live cells
LSM
Tu/Epi
(debris removal)
B
CD45+ purified
LNP
LNP-mIFNa2
Supernatant (CD45-enriched)
TU-LNP
TU- LNP-mIFNα2
IgG-APC
PD-L1-APC
CD45-PE
C
UMAP_1
UMAP_2
E
Activated NK cells
Basophils
CD8 Tex-intermediate
cDC1
cDC2
DCs
Endothelial Cells
eTregs
Fibroblasts
Fibroblasts (Pi15+)
IgD+CD20+ Naive Mature B cells
IL-17-producing gd T cells
Ly6+ Inflammatory Monocytes
Macrophages
Mature Naive/Follicular B cells
Naive B cells
Naive/Stem-like CD8 T cells (Tpex-early)
pDCs
Plasma Cells
Proliferating Cytotoxic NK cells
Tgd cells
Tpex-like T cells
Tumor-Associated Macrophages
Tumor-Associated Neutrophils
Tumor/Epithelial
D
Features
Identity
Percent Expressed
Average Expression
F
Features
Identity
Percent Expressed
Average Expression

**Figure S5. scRNA-seq clustering and cell type annotation of immune cells in lung metastases. A.** Experimental scheme of $CD45^+$ and $CD45^+$ cell-depleted cell populations for scRNAseq. **B**. Dot plots of flow cytometry of purity of purified $CD45^+$ cells (left) and residual $CD45^-$ cell-enriched populations. **C**. UMAP plot of all sequenced cells colored by unsupervised cluster identity (clusters 0-30). **D.** Dot plot of marker gene expression used for cluster identity assignments (clusters 0-30 on x-axis, feature genes on y-axis). Dot size indicates the percentage of cells expressing the gene; color indicates average scaled expression. **E.** UMAP plot showing annotated cell type identities. **F.** Dot plot of marker gene expression confirming annotated cell type identities across all identified populations.

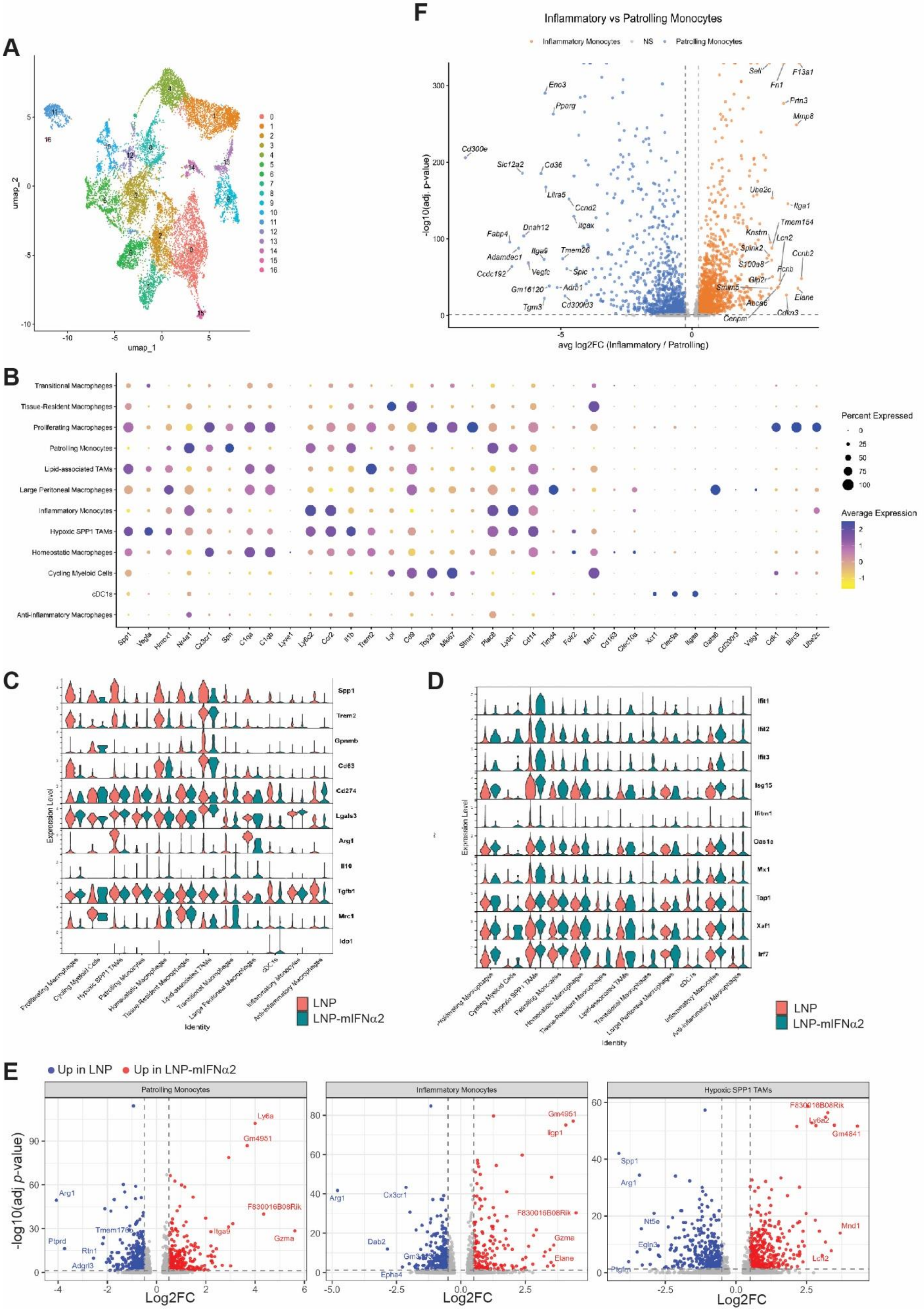
A
B
C
D
E
F
Inflammatory vs Patrolling Monocytes
Inflammatory Monocytes
NS
Patrolling Monocytes
avg log2FC (Inflammatory / Patrolling)
-log10(adj. p-value)
Percent Expressed
Average Expression
LNP
LNP-mIFNα2
Up in LNP
Up in LNP-mIFNα2
Patrolling Monocytes
Inflammatory Monocytes
Hypoxic SPP1 TAMs
Log2FC
-log10(adj p-value)
Expression Level
Identity
umap_1
umap_2

**Figure S6. Subclustering of myeloid cells reveals LNP-mIFNα2-driven elimination of suppressive macrophage states and emergence of IFN-responsive monocytes. A.** UMAP of subclustered monocyte and macrophage populations, colored by unsupervised cluster identity. **B.** Dot plot of key marker genes used for myeloid cell type annotations. Dot size indicates percent of cells expressing the gene; color indicates average scaled expression. **C.** Violin plots of immunosuppressive marker gene expression across myeloid subtypes, split by sample (LNP: pink; LNP-mIFNα2: green). **D.** Violin plots of IFN-response marker gene expression across myeloid subtypes, split by sample (LNP: pink; LNP-mIFNα2: blue). **E.** Volcano plots showing differentially expressed genes in the indicated re-clustered myeloid populations (patrolling monocytes, inflammatory monocytes, and hypoxic $SPP1^{+}$ tumor-associated macrophages) comparing LNP control and LNP-mIFNα2-treated tumors. Red, genes upregulated in LNP-mIFNα2; blue, genes upregulated in LNP control. **F.** Volcano plot of differentially expressed genes comparing inflammatory monocytes versus patrolling monocytes clusters. Orange=enriched in inflammatory monocytes; blue=enriched in patrolling monocytes.

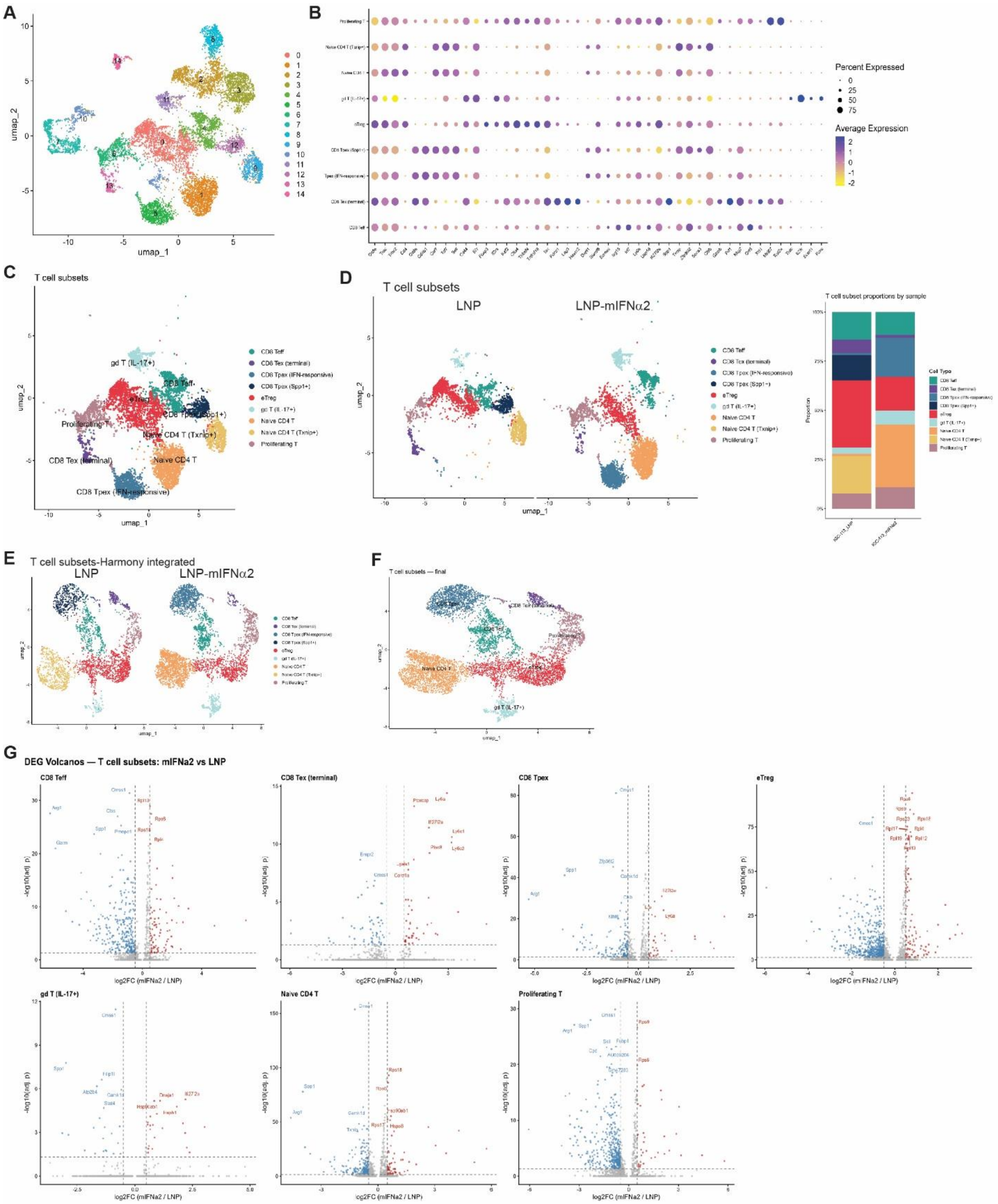

**Figure S7. Subclustering of T cells reveals LNP-mIFNα2-driven expansion of progenitor-exhausted CD8 T cells and contraction of effector Tregs. A.** UMAP of subclustered T cell populations, colored by unsupervised cluster identity. **B.** Dot plot of marker genes used for T cell subtype annotations. **C-D.** UMAP of annotated T cell subsets (C), split by sample (LNP, LNP-mIFNα2, D), and stacked bar chart of subset proportions by sample (D).

**E.** Harmony-integrated UMAP of annotated T cell subsets, split by sample. **F.** UMAP of T cell subsets following collapse of the CD8 Tpex and Naive CD4 T clusters, yielding the final seven-subset annotation scheme. **G.** Volcano plots of differentially expressed genes (LNP vs LNP-mIFNα2) within each T cell subset: CD8 Teff, CD8 Tex-term, CD8 Tpex, eTreg, γδ T (IL-17$^{+}$), Naive CD4 T, and Proliferating T cells. Blue: upregulated in LNP. Red: upregulated in LNP-mIFNα2.

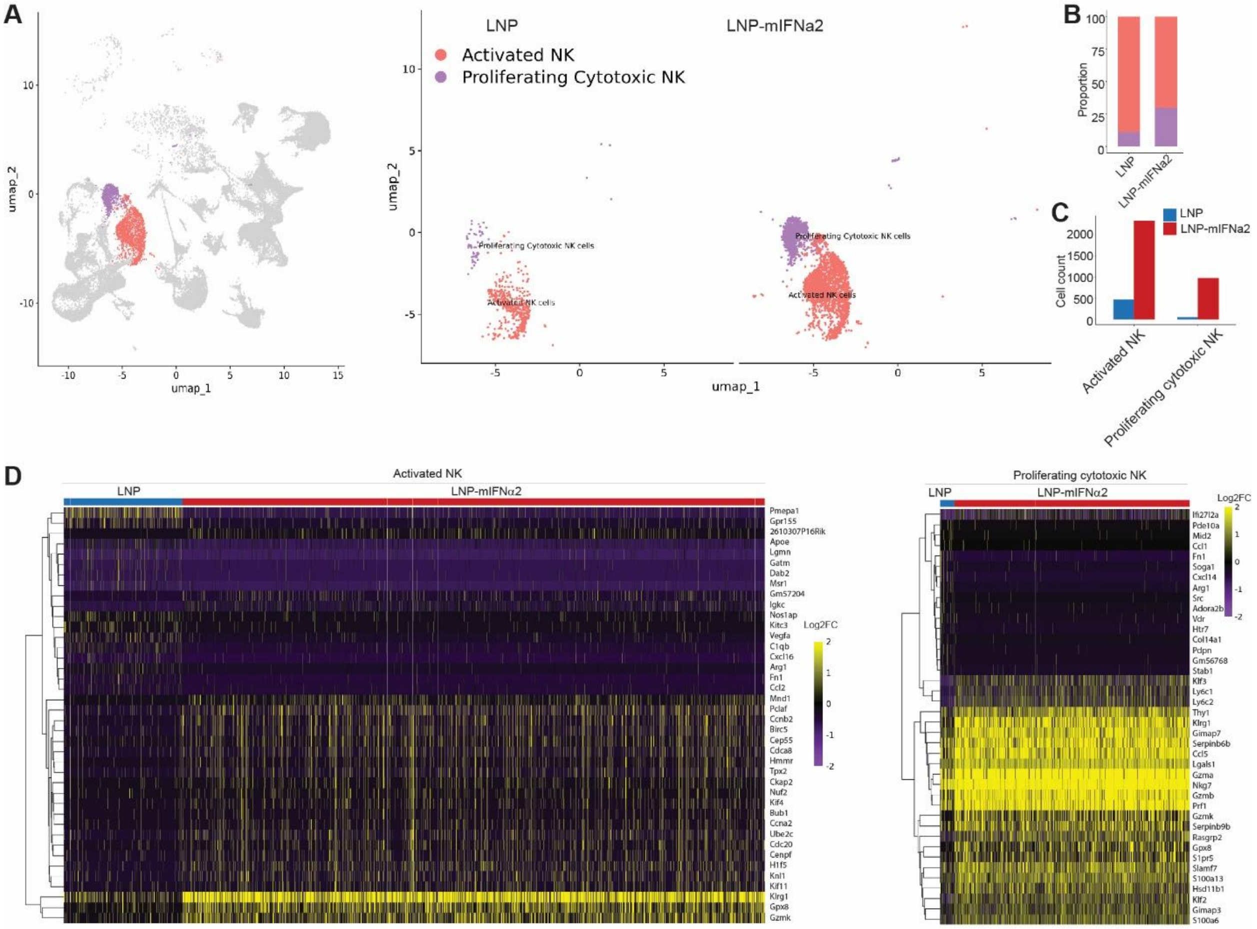


**Figure S8. LNP-mIFNα2 expands and transcriptionally reprograms intratumoral NK cell subsets. A.** UMAP plots showing activated NK cells and proliferating cytotoxic NK cells from LNP- and LNP-mIFNα2-treated tumors (left, combined; right, split by treatment condition). **B.** Stacked bar chart showing the proportional composition of activated NK and proliferating cytotoxic NK cells within LNP- and LNP-mIFNα2-treated samples. **C.** Bar chart showing absolute cell counts of activated NK and proliferating cytotoxic NK cells per condition. **D.** Heatmaps of differentially expressed genes (log2 fold-change) for activated NK cells (left) and proliferating cytotoxic NK cells (right), comparing LNP to LNP-mIFNα2. Each row is a gene; each column is a cell.

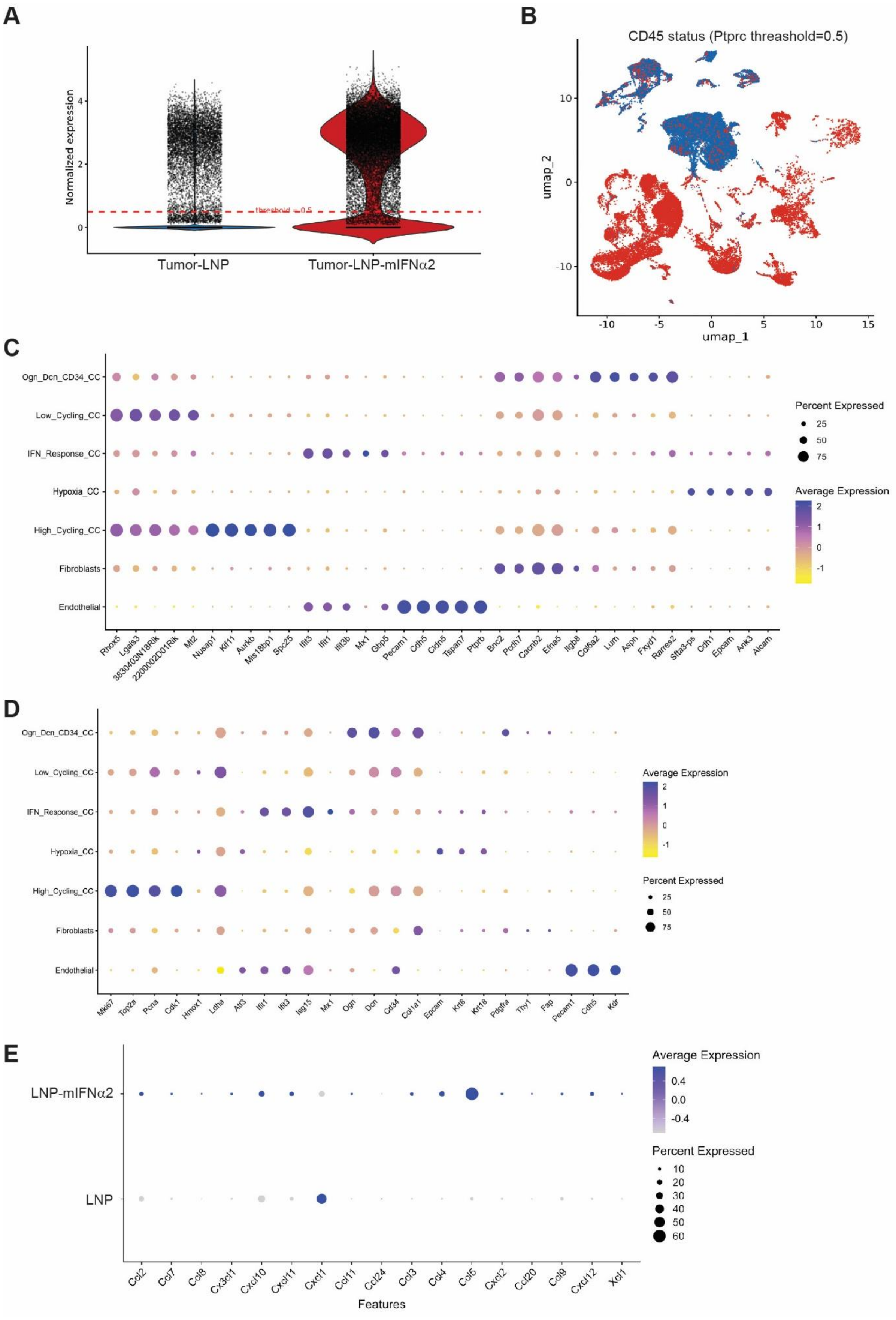

A
Normalized expression
threshold = 0.5
Tumor-LNP
Tumor-LNP-mIFNα2
B
CD45 status (Ptprc threashold=0.5)
umap_1
umap_2
C
Ogn_Dcn_CD34_CC
Low_Cycling_CC
IFN_Response_CC
Hypoxia_CC
High_Cycling_CC
Fibroblasts
Endothelial
Rhox5 Lgals3 3830403N18Rik 2200002D01Rik Mt2 Nusap1 Kif11 Aurkb Mis18bp1 Spc25 Ifit3 Ifit1 Ifit3b Mx1 Gbp5 Pecam1 Cdh5 Cldn5 Tspan7 Ptprb Bnc2 Pcdh7 Cacnb2 Efna5 Itgb8 Col6a2 Lum Aspn Fxyd1 Rarres2 Sfta3-ps Cdh1 Epcam Ank3 Alcam
Percent Expressed
Average Expression
D
Ogn_Dcn_CD34_CC
Low_Cycling_CC
IFN_Response_CC
Hypoxia_CC
High_Cycling_CC
Fibroblasts
Endothelial
Mki67 Top2a Pcna Cdk1 Hmox1 Ldha Atf3 Ifit1 Ifit3 Isg15 Mx1 Ogn Dcn Cd34 Col1a1 Epcam Krt8 Krt18 Pdgfra Thy1 Fap Pecam1 Cdh5 Kdr
Average Expression
Percent Expressed
E
LNP-mIFNα2
LNP
Ccl2 Ccl7 Ccl8 Cx3cl1 Cxcl10 Cxcl11 Cxcl1 Ccl11 Ccl24 Ccl3 Ccl4 Ccl5 Cxcl2 Ccl20 Ccl9 Cxcl12 Xcl1
Features
Average Expression
Percent Expressed

**Figure S9. Transcriptional annotation of the non-immune (CD45$^-$) tumor and stromal compartment in lung metastases. A.** Violin plot of *Ptprc* (CD45) normalized expression in LNP and LNP-mIFNα2-treated tumor samples. Dashed red line indicates the threshold used to classify cells as CD45$^-$ (tumor/stromal cells) versus CD45$^+$ (immune cells) for downstream filtering. **B.** UMAP of the integrated scRNA-seq dataset, colored by CD45 status based on the *Ptprc* threshold defined in (A). Blue: CD45$^-$ cells; Red: CD45$^+$ cells. **C.** Dot plot of the top 5 differentially expressed genes (by log2 fold-change, ranked by pct.1-pct.2) across the annotated CD45$^-$ cell subtypes as indicated. **D.** Dot plot of canonical marker gene expression confirming annotation of the seven CD45$^-$ cell subtypes shown in (C). **E.** Dot plot of chemokine gene expression restricted to tumor cells only, split by sample (LNP vs LNP-mIFNα2).
